# Water levels primarily drive variation in photosynthesis and nutrient use of scrub Red Mangroves in the southeastern Florida Everglades

**DOI:** 10.1101/2021.01.27.428494

**Authors:** J. Aaron Hogan, Edward Castañeda-Moya, Lukas Lamb-Wotton, Tiffany Troxler, Christopher Baraloto

**Affiliations:** Department of Biological Sciences, Florida International University, 11200 SW 8th Street, OE-167, Miami, FL USA 33199; Institute of Environment, Florida International University, 11200 SW 8th Street, OE-148, Miami, FL USA, 33199; Department of Earth and Environment, Florida International University, 11200 SW 8th Street, AHC5-360, Miami, FL, USA 33199

**Keywords:** Scrub mangroves, Florida Coastal Everglades, photosynthesis, porewater salinity, water levels, inundation, *Rhizophora mangle*

## Abstract

We investigated how mangrove-island micro-elevation (i.e., habitat: center vs. edge) affects tree physiology in a scrub mangrove forest of the southeastern Everglades. We measured leaf gas exchange rates of scrub *Rhizophora mangle* L. trees monthly during 2019, hypothesizing that CO_2_ assimilation (*A_net_*) and stomatal conductance (*g_sw_*) would decline with increasing water levels and salinity, expecting more-considerable differences at mangrove-island edges than centers, where physiological stress is greatest. Water levels varied between 0 and 60 cm from the soil surface, rising during the wet season (May-October) relative to the dry season (November-April). Porewater salinity ranged from 15 to 30 ppt, being higher at mangrove-island edges than centers. *A_net_* maximized at 15.1 µmol m^-2^ s^-1,^ and *g_sw_* was typically <0.2 mol m^-2^ s^-1^, both of which were greater in the dry than the wet season and greater at island centers than edges, with seasonal variability being roughly equal to variation between habitats. After accounting for season and habitat, water level positively affected *A_net_* in both seasons but did not affect *g_sw_*. Our findings suggest that inundation stress (i.e., water level) is the primary driver of variation in leaf gas exchange rates of scrub mangroves in the Florida Everglades, while also constraining *A_net_* more than *g_sw_*. The interaction between inundation stress due to permanent flooding and habitat varies with season as physiological stress is alleviated at higher-elevation mangrove-island center habitats during the dry season. Freshwater inflows during the wet season, increase water levels and inundation stress at higher-elevation mangrove-island centers, but also potentially alleviate salt and sulfide stress in soils. Thus, habitat heterogeneity leads to differences in nutrient and water acquisition and use between trees growing in island centers versus edges, creating distinct physiological controls on photosynthesis, which likely affect carbon flux dynamics of scrub mangroves in the Everglades.

## INTRODUCTION

Global climate change is affecting coastal mangrove ecosystems unprecedentedly, principally through increased flooding and saltwater intrusion (Pezeshki et al. 1990a, Yu et al. 2019). Increases in flooding severity and salinity due to sea-level rise (SLR) have the potential to push ecosystems to degraded alternative stable states, where biogeochemical cycles (e.g., carbon sequestration and storage potential) are impaired (Neubauer et al. 2013, Tully et al. 2019, Yu et al. 2019). Mangrove wetlands are particularly susceptible to SLR because of their position between terrestrial and marine ecosystems (Field 1995, Ellison and Farnsworth 1997). Mangrove species have developed considerable variation in crucial life-history traits, such as rates of photosynthesis, water- and nutrient-use efficiencies, growth rates, and biomass allocation ratios in response to the interactions among resources (e.g., light and nutrients), regulators (e.g., salinity, sulfides), and inundation – Twilley and Rivera-Monroy 2005, Alongi 2008, Twilley and Rivera-Monroy 2009, Castañeda-Moya et al. 2013). Due to such physiological flexibility and the significant carbon sequestration and storage capacity of mangroves across a variety of geomorphic settings (e.g., karstic vs. deltaic; Mcleod et al. 2011, Murdiyarso et al. 2015, Lovelock et al. 2017, Rovai et al. 2018), there is an increasing need to strengthen our understanding of the effects of SLR and saltwater intrusion on mangrove tree physiology to assess trajectories of ecosystem structure and function in response to global change drivers.

Scrub mangrove forests, dominated by *Rhizophora mangle* L., are typical in Caribbean karstic environments (Cintron et al. 1978, Lugo and Snedaker 1974). The stunted physiognomy (i.e., reduced growth and development) of scrub mangroves results from severe nutrient (e.g., phosphorus, P) limitation, prolonged or permanent inundation with little tidal influence, and seasonal water stress (Feller 1995, Koch and Snedaker 1997, Cheeseman and Lovelock 2004, Medina et al. 2010). Scrub mangrove forests develop distinct landscape patterning, forming mangrove-island clusters with higher-elevations than their surrounding shallow open-water ponds and channels (Figure 1A). Soil elevation gradients result from differences in root biomass stocks and production, leaf litter accretion, and wood deposition (McKee et al. 2007, McKee 2011, Krauss et al. 2014). For example, in scrub mangrove-islands of the southeastern Florida Everglades, island center habitats have 66% more root biomass and 52% more root production than island edges (Castañeda-Moya et al. 2011), which leads to spatial differences in soil elevation among island habitats. These differences in soil elevation interact with environmental gradients (e.g., hydroperiod, salinity) along the intertidal zone in complex ways to affect mangrove physiology (e.g., rates of net CO_2_ assimilation – *A_net_*, growth rates, or sap flux) at variable scales (Medina and Francisco 1997, Twilley et al. 1998, Medina et al. 2010, Twilley et al. 2017).

**Figure 1.**
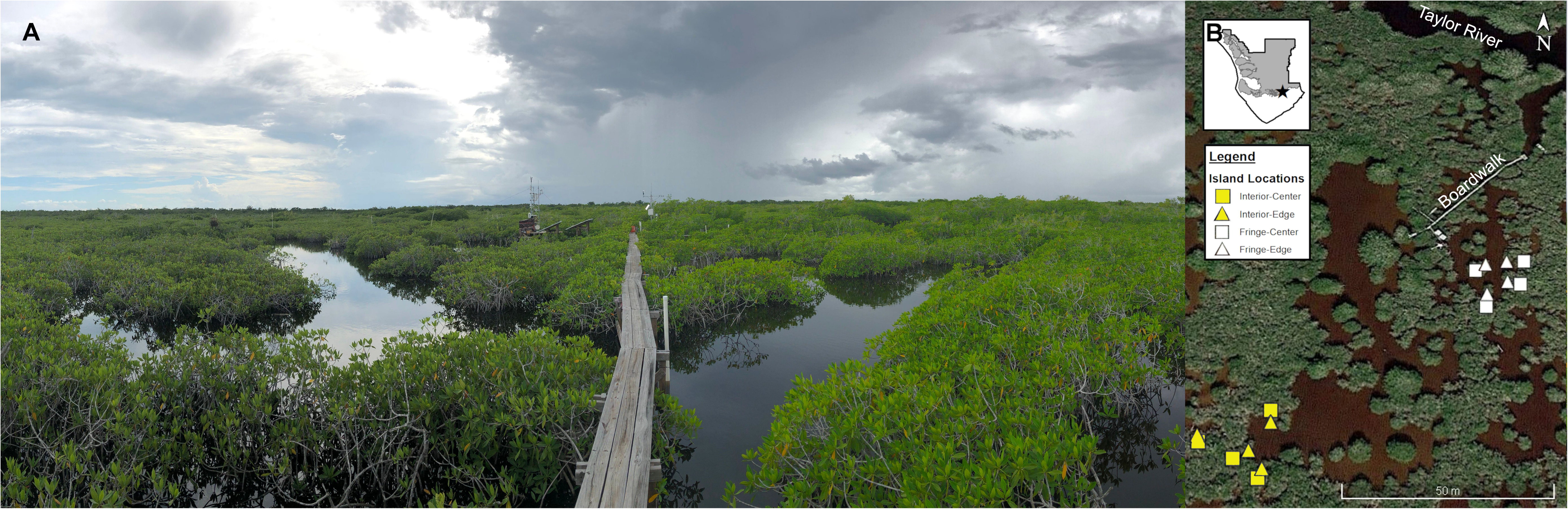
**A)** Photograph of TS/Ph-7 shows scrub R. mangle tree islands that characterize the study site. Mangrove canopy heights are approximately 1.5-2 m tall, facilitating canopy measurements of leaf physiology. Boardwalk (1.3 m height) is pictured for reference. **B)** Aerial view (Google Earth) of mangrove-islands measured for this study within TS/Ph-7, near the mouth of the Taylor River in southeastern Florida Coastal Everglades, USA. The inset shows the location of TS/Ph-7 within the boundary of Everglades National Park. Colors indicate scrub mangroves and fringe (white) and interior zones (yellow) relative to Taylor River. Symbols denoted paired higher-elevation center and lower-elevation edge habitats for each mangrove-island (squares and triangles, respectively).

Hydrological dynamics (e.g., depth and duration of inundation) can cause mangrove physiological stress, which cascades to affect carbon cycle dynamics and other biogeochemical processes across spatial and temporal scales (Medina 1999, Castañeda-Moya et al. 2013, Twilley et al. 2017, 2019). Although mangrove species can tolerate flooded conditions, they are still susceptible to damage if plants become entirely submerged for days to weeks (Wanless 1998, Mendelssohn and McKee 2000, McKee 2011). Inundation stress typically decreases rates of leaf gas exchange (e.g., *A_net_*, transpiration) and tree growth in mangroves (He et al. 2007, Cardona-Olarte et al. 2013). For example, greenhouse studies have revealed a 20% reduction in maximum *A_net_* when mangrove seedlings and saplings were subjected to short-term intermittent seawater flooding (6 to 22 days, Krauss et al. 2006). Mangrove leaf gas exchange is further affected by how seawater flooding interacts with fresh water and nutrient inputs (Wolanski 1992). For instance, a significant reduction in stomatal conductance (*g_sw_*) and leaf water potential in *Bruguiera gymnorrhiza* (L.) Lam. seedlings occurred when exposed to prolonged flooding for up to 80 days with 33% seawater compared to the control plants; however, seedlings flooded with fresh water for 80 days showed an increase in both parameters (Naidoo 1983). In contrast, seedlings of *Avicennia germinans* (L.) L. and *Laguncularia racemosa* (L.) C.F.Gaertn. exposed to permanent flooding with 23% seawater showed no change in *g_sw_*, *A_net_*, or intrinsic water use efficiency (*wue*) but had reduced leaf area (Krauss et al. 2006). Hydrologic conditions can further negatively influence mangrove physiology through the interaction with soil phytotoxins (i.e., sulfides), produced as by-products of low oxygen availability and soil redox conditions due to permanent flooding, which can potentially depress water and nutrient uptake and affect rates of leaf gas exchange (Nickerson and Thibodeau 1985, McKee 1993, Ball 1996, Pezeshki and DeLaune 2012, Lamers et al. 2013). Regarding permanently inundated scrub mangroves, such as those in the southeastern Florida Everglades, how seasonal dynamics interact with inundation levels to influence leaf and forest carbon uptake dynamics is not entirely understood.

Mangroves are highly adapted to tolerate salt stress, yet salinity has the most significant impact on forest productivity, tree growth rates, and rates of leaf gas exchange. The adverse effects of increasing salinity are most evident along steep salinity gradients (i.e., those >30 ppt), particularly in dry environments (Lugo and Snedaker 1974, Cintron et al. 1978, Medina and Francisco 1997, Reef and Lovelock 2015). Salt stress variably affects mangrove tree physiology, depending on species-specific salt tolerance levels and mechanisms for processing salt (Parida and Jha 2010, Reef and Lovelock 2015). For example, *R. mangle* naturally inhabits Neotropical environments with salinities from near zero to around 35 ppt but may also be found in dry coastal environments with salinities up to 50-60 ppt (Cintron et al. 1978, Cardona-Olarte et al. 2006). All mangroves can exclude salt through the roots; however, *R. mangle* is a highly efficient salt excluder because its roots essentially prevent salt from entering the plant. Additionally, *R. mangle* lacks the excretory glands that other mangrove species (e.g., *L. racemosa*) use to excrete salt once it has entered the plant. As such, the xylem of *R. mangle* is 100 times less saline than seawater (Scholander et al. 1962, Scholander 1968, Medina and Francisco 1997, Tomlinson 2016) because of the Casparian strip (Lawton et al. 1981) and ultrafiltration by cell membranes in the thick aerenchyma and cortical layers of its root tissues (Field 1984, Werner and Stelzer 1990). However, some salt still enters the plant through the roots, which has a deleterious effect on the physiology of *Rhizophora* trees, causing decreases in growth and *A_net_* rates, and water and nutrient use efficiencies (Ball 1988, Clough and Sim 1989, Lugo et al. 2007, Medina et al. 2010, Cardona-Olarte et al. 2013).

Mangrove *A_net_* varies widely with the environment (e.g., water and salinity levels) and nutrient availability. *A_net_* for *R. mangle* maximizes around 20 µmol m^-2^ s^-1^ (Golley et al. 1962, Bjorkman et al. 1988, Lin and Sternberg 1992, Lovelock and Feller 2003, Lugo et al. 2007, Ball 2009); however, *A_net_* for scrub mangroves is lower, generally ranging from <5 µmol m^-2^ s^-1^ (Golley et al. 1962, Cheeseman et al. 1997, Cheeseman and Lovelock 2004) to roughly 13 µmol m^-2^ s^-1^ (Lugo et al. 2007, Barr et al. 2009). A field study from Jobos Bay in southern Puerto Rico demonstrated a significant decrease in *R. mangle A_net_* (from 12.7 to 7.9 µmol m^-2^ s^-1^) and *g_sw_* (from 0.28 to 0.19 mol m^-2^ s^-1^) when comparing fringe habitats at 35 ppt salinity to inland salt flat habitats at 80 ppt (Lugo et al. 2007). Reductions in *A_net_* and *g_sw_* were accompanied by changes in leaf morphology (i.e., smaller specific leaf area, SLA), reduced nutrient-use efficiency, and increased nutrient resorption, demonstrating how environmental effects on mangrove physiology can have consequences for within plant nutrient dynamics, and thus ecosystem functioning. Therefore, increasing salinity decreases *A_net_* and *g_sw_* and increases intrinsic water use efficiency (*wue*, defined as *A_net_/g_sw_*) in mangroves, with *Rhizophora* species exemplifying these trends (Ball 2009, Clough and Sim 1989). Moreover, the high salt tolerance of mangrove species leads to strong stomatal control, which creates dynamics between *A_net_* and water use which depend on relative reduction in transpiration rates versus the degree to which leaves are biochemically limited to fix carbon (e.g., via RUBISCO carboxylation efficiency versus RUBP regeneration) at low stomatal conductance (Sobrado 2000, Lovelock and Feller 2003, Ball 2009). For instance, *R. mangle* has succulent leaves with lower *wue* than more salt-tolerant species (i.e., *A. germinans* or *L. racemosa*); however, *R. mangle* has greater water transport efficiency in stems than more salt-tolerant species (Sobrado 2000). Thus, when considering the effects of salinity on leaf gas exchange rates, plant water use must be considered in concert because both *A_net_* and *g_sw_* decline similarly with increasing salinity, effectively creating co-limitation of photosynthesis at moderate to high salinities (Ball 2009).

In mangrove wetlands of the Florida Everglades, variation in environmental gradients, including hydroperiod (e.g., duration of inundation) and soil P availability, control mangrove forest structure and function (e.g., biomass and litterfall production) across the coastal landscape (Chen and Twilley 1999; Castañeda-Moya et al. 2011, 2013). Yet, how the interaction between water level dynamics and salinity affects mangrove leaf gas exchange rates *in situ* is not entirely understood. Experimental evidence using *R. mangle* seedlings from south Florida showed that inundation created a greater degree of physiological stress than salinity levels; however, salinity accelerated the adverse effects of inundation stress on leaf function over time (Cardona-Olarte et al. 2013). In contrast, other studies have reported no apparent effect of water levels or flooding duration on rates of mangrove gas exchange, although inundation duration decreased variability in leaf gas exchange measurements (Hoppe-Speer et al. 2011). Using Florida mangroves, Krauss et al. (2006) found that short-term intermittent flooding decreased rates of leaf gas exchange relative to unflooded or permanently flooded greenhouse-grown seedlings, but that for *in situ* established *R. mangle* saplings growing along a natural tidal inundation gradient along Shark River in the southwestern Everglades, flooding led to increases in *A_net_* and *wue*. Permanent flooding leads to decreases in *A_net_* and *g_sw_* rates in most wetland plants (Kozlowski 1997); however, how inundation dynamics interact with salinity along the intertidal zone to influence mangrove physiology at different spatial and temporal scales in south Florida mangroves remains largely unknown. Further, global change-driven SLR and saltwater intrusion in South Florida coupled to large-scale freshwater diversion have accelerated mangrove encroachment into inland freshwater wetlands over the past 60 years (Ross et al. 2000). As SLR continues, it is imperative to quantify the relative effects of inundation and salinity on mangrove physiology and subsequent ecosystem functioning (e.g., carbon flux) in the region.

Here, we present a comprehensive, one-year analysis of the seasonal effects of salinity (surface and porewater) and water levels on leaf gas exchange rates of *R. mangle* scrub mangroves in southeastern Florida Everglades. We focused our sampling on mangrove-islands with noticeable micro-elevational differences to understand the magnitude of influence of water levels and salinity on *R. mangle* tree physiology between mangrove-island center and edge habitats. We addressed the following questions: (1) how do rates of leaf gas exchange (e.g., *A_net_*, *g_sw_*) vary with mangrove-island micro-elevation (center vs. edge habitats)? (2) how does leaf gas exchange respond to seasonal changes in salinity and water levels? (3) how do water- and nutrient-use efficiencies of *R. mangle* leaves vary between mangrove-island center and edge habitats? We hypothesized that *A_net_* would be greater for *R. mangle* leaves in higher-elevation center habitats than lower-elevation edges. We also expected that *A_net_* should vary little with season and that seasonal variation in *g_sw_* would be less than variation in *A_net_*, relative to the range of variability among leaves because of strong control on *g_sw_* and potential for decoupling to some degree between *g_sw_* and *A_net_.* Moreover, given that scrub mangroves in Taylor River basin are strongly limited by phosphorus (i.e., soil N:P = 102-109 – Castañeda-Moya et al. 2013), *R. mangle* plants should have high rates of P resorption. Finally, we predicted that trees within mangrove-island centers function at a higher physiological level (i.e., with greater rates of *A_net_* and less of a relative reduction in *g_sw_*) due to lower levels of inundation and salt stress, and should, therefore, have greater *wue* (Ball 2009) and higher relative rates of nutrient resorption (Lugo et al. 2007, Medina et al. 2010) than trees at island edges. However, the magnitude of the reduction in *g_sw_* relative to *A_net_*, because of inundation stress at mangrove-island edge habitats, should drive patters in *wue*.

## METHODS

### Study site

This study was conducted in the southeastern region of Everglades National Park in a mangrove site known as Taylor Slough/Panhandle-7 (TS/Ph-7: 25.197°N, 80.642°W, Figure 1B), one of the six mangrove sites established in 2000 as part of the Florida Coastal Everglades Long-Term Ecological Research (FCE-LTER) program (Childers 2006; http://fcelter.fiu.edu). TS/Ph-7 is located approximately 1.5 km inland from Florida Bay in the downstream section of the Taylor River. Mangroves zones at TS/Ph-7 are dominated by scrub *R. mangle* L. trees, with clusters of *L. racemosa* L. and *Conocarpus erectus* L. – a mangrove associate, intermixed with low densities of freshwater grasses *Cladium jamaicense* (Crantz) Kük and *Eleocharis cellulosa* Torr. (Loveless 1959). Mangrove tree heights reach 1.5 to 2 m (Ewe et al. 2006, Figure 1A).

The substrate at this site is organic mangrove peat soil (∼1 m depth) overlying the karstic bedrock (depth ∼1.5-2 m, Table 1; Castañeda-Moya et al. 2011, Ewe et al. 2006). Surface (0-45 cm depth) soils at TS/Ph-7 have high organic matter content (71%), low bulk density (0.16 g cm^-3^), low total nitrogen (TN, 2.5 mg cm^-3^), and low total phosphorus (TP, 0.06 mg cm^-3^) concentrations, resulting in a highly P-limited environment with soil N:P ratios of about 102 (Castañeda-Moya et al. 2013). Mangrove zones in Taylor River are permanently flooded for most of the year, with an annual flooding duration averaging 360 d yr^-1^ from 2001 to 2005. The permanent flooding results in anoxic soil conditions and buildup of porewater sulfide (range: 0.5-2 mM) throughout the year that constrains mangrove growth (Castañeda-Moya et al. 2011, 2013). The tidal effect is negligible in Taylor River, and water flow and hydrology are determined by seasonal precipitation, upland runoff, and wind (Michot et al. 2011, Sutula et al. 2001). The interaction between low P fertility and permanent flooding conditions results in the formation of scrub forests with restricted tree height and low aboveground productivity, high root biomass allocation and high root: shoot ratios compared with riverine mangrove forests along Shark River estuary in southwestern FCE (Ewe et al. 2006, Castañeda-Moya et al. 2011, 2013).

**Table 1.** Mean (± 1 SE) soil surface elevation, bedrock elevation, and soil depth, for open water, mangrove-island edge and center habitats in the fringe and interior scrub mangrove areas at TS/Ph-7 in southeastern Florida Coastal Everglades. Elevation measurements are referenced to the North American Vertical Datum 1988 (NAVD88). Letters denote statistically different groupings via Tukey HSD post-hoc test (p < .05).

| Mangrove Location | Habitat | Soil Surface Elevation (NAVD88, m) | Bedrock Elevation (NAVD88, m) | Soil depth (m) |
| --- | --- | --- | --- | --- |
| Fringe | Open water | $-0.63 \pm 0.05^C$ | $-1.84 \pm 0.05^A$ | $1.22 \pm 0.05^C$ |
| | Island edge | $-0.41 \pm 0.03^B$ | $-1.83 \pm 0.02^A$ | $1.43 \pm 0.05^{BC}$ |
| | Island center | $-0.14 \pm 0.02^A$ | $-1.86 \pm 0.06^A$ | $1.72 \pm 0.05^A$ |
| Interior | Open water | $-0.84 \pm 0.02^D$ | $-2.01 \pm 0.15^A$ | $1.18 \pm 0.14^C$ |
| | Island edge | $-0.49 \pm 0.03^B$ | $-1.78 \pm 0.04^A$ | $1.39 \pm 0.04^C$ |
| | Island center | $-0.15 \pm 0.02^A$ | $-1.77 \pm 0.03^A$ | $1.62 \pm 0.03^{AB}$ |

South Florida has a subtropical savanna climate per the Köppen climate classification, where the average air temperature is between 20 and 30°C and relative humidity is high (70-80%). Rainfall and evapotranspiration vary interannually and average 1500 and 1300 mm year^-1^, respectively (Abiy et al. 2019). In the Everglades, 60% of the precipitation occurs during the wet season, and only 25% during the dry season (Duever et al. 1994). Analysis of long-term (110-year) rainfall trends for South Florida has shown that the annual hydrologic regime can be divided into two seasons: a wet season from May to October and a dry season from November to April (Abiy et al. 2019). For the 2019 calendar year, temperature and relative humidity data were collected from an eddy covariance flux tower installed at TS/Ph-7. Rainfall data were collected from a nearby meteorological station (station name: “Taylor_River_at_mouth”) managed by the US Geological Survey as a part of the Everglades Depth Estimation Network (https://sofia.usgs.gov/eden).

### Experimental design

Eight distinct mangrove-islands of similar size (3-5 m in diameter) were selected for repeated measurements of leaf photosynthesis and physicochemical variables from January to December 2019. Mangrove-islands were selected within previously-established permanent vegetation plots (two 20×20 m plots) based on their location relative to the shoreline (i.e., Taylor River), with four islands located in the fringe mangrove zone (∼50-60 m from the edge) and four islands located in the interior forest (∼100-110 m inland; Figure 1B). Mangrove-islands with distinct micro-elevational gradients were selected, having higher soil elevation center habitats and lower-elevation edge habitats. Mangrove-islands are surrounded by open water ponds (Figure 1A) and remain flooded for most of the year, except the center island habitats during the dry season (Castañeda-Moya et al. 2011, 2013, Figure S1).

Within each mangrove-island, a higher-elevation center and a lower-elevation edge habitat were each permanently marked with an aluminum rod. At these locations, soil surface elevation was measured for all mangrove-islands at both habitats, in addition to six measurements in the adjacent shallow ponds surrounding mangrove-islands. Measurements were taken using real-time kinematics referenced to the 1988 North American Vertical Datum (NAVD88) with a Trimble R8 global navigation satellite system receiver (Trimble; Sunnyvale, CA, USA), which has a horizontal accuracy of ±1 cm and vertical accuracy of ±2 cm.

### Water level and salinity measurements

Water levels relative to the soil surface were measured monthly with a meter stick at the permanent aluminum rods established at all island habitats. Continuous measurements of water levels relative to the soil surface were recorded for the duration of the 2019 calendar year (see Figure S2 for details). Continuous data were used to confirm trends in water levels and porewater salinity measurements made by hand across islands. We use the measurements taken by hand at each island as the predictors in our models of leaf gas exchange. A porewater sample was collected at 30 cm depth at each habitat using a 60 ml syringe attached to a stopcock and a rigid tubing probe (3/16” Ø). Porewater temperature and salinity were measured using a handheld YSI conductivity-salinity-temperature meter (model Pro 30, YSI Inc., Yellow Springs, OH, USA). A sample of surface water (when present) was also collected at each island habitat to measure salinity and temperature.

### Leaf gas exchange measurements

Photosynthetic gas exchange measurements of *R. mangle* leaves were conducted once a month (9:00 AM to 1:00 PM, typically during sunny days) at eight scrub mangrove-islands from January to December 2019 using a Li-COR Li-6800 portable photosynthesis system (Li-COR Inc., Lincoln, NE, USA). At each island habitat (center vs. edge), five mature green leaves were randomly selected from top mangrove branches. Fully developed and healthy (i.e., without herbivory) green leaves from the second-most distal pair of leaves were chosen. The Li-6800 was clamped onto each leaf and held until machine stability was reached (which typically happened in 2-3 minutes), wherein data points were logged.

The environmental configuration of the Li-6800 was: flow rate of 600 µmol s^-1^, 50-70% relative humidity of the incoming air (slightly drier than ambient air to prevent condensation in the instrument), 400 µmol mol^-1^ CO_2_ concentration, and light level of 1000 µmol m^-2^ s^-1^, which was determined to be non-limiting and similar to ambient environmental conditions. We used five stability criteria, which were all assessed over a 15 s interval: the slope of *A_net_* being <1 µmol m^-2^ s^-1^, the slope of the concentration of intracellular CO_2_ (*c_i_*, which is a calculated parameter using the difference in CO_2_ concentrations between IRGAs) being <5 µmol mol^-1^, the slope of *g_sw_* being < 0.5 mol m^-2^ s^-1^, the slope of the transpiration rate (*E*) being <1 mol m^-2^ s^-1^, and the slope of the difference in air-water vapor concentration between the sample and reference IRGA (*ΔH_2_O*) being <1 mmol mol^-1^. All five stability criteria were met before logging data. Air temperature within the leaf chamber was not controlled but allowed to vary with the ambient conditions at the site, ranging from 26.1 to 32.0°C. Thus, leaf temperatures ranged from 25.85 - 32.44 °C, averaging 29.41 ± 0.08 °C, in the wet season, and ranged from 25.06 - 28.72 °C, averaging 26.79 ± 0.04 °C, in the dry season. We calculated intrinsic *wue* as the ratio of leaf net CO_2_ uptake to leaf gas exchange (i.e., *A_net_*/*g_sw_*/1000, where we divide by 1000 to get *wue* in mmol mol^-1^).

### Measurement of leaf functional traits, nutrient content, and isotopic signatures

During the monthly photosynthesis measurements in February, May, August, and November measured mature green leaves (n = 5 per habitat, 40 in total) were collected at half of the islands (four of the eight islands with two per location) for determination of leaf functional traits and total carbon (TC), nitrogen (TN), and phosphorus (TP) content. Leaves were numbered, placed in a sealed, moist bag to prevent water loss, and transported to the laboratory in a cooler with ice for further analyses. Five senescent (i.e., yellowing) leaves were also collected from trees in the same islands at the same time to determine carbon and nutrient content. Leaves were removed from bags, wiped dry, and immediately weighed to obtain leaf fresh mass at the laboratory. Green leaves were then scanned at high resolution and oven-dried for at least 72 hours at 60°C to constant weight before recording their dry mass. Leaf area was measured using ImageJ (Schneider et al. 2012). Leaf dry mass was recorded and used to calculate leaf dry matter content (LDMC) as the ratio of the dry leaf mass (in mg) to its fresh mass (in g, mg g^-1^), percent leaf water content (1000-LDMC; %), and SLA, the ratio of leaf dry weight to leaf area (cm^2^ g^-1^). These methods followed Cornelissen et al. (2003).

For nutrient analyses, composite leaf samples containing the five leaves from each island habitat per collection were ground into a fine powder using a vibrating ball mill (Pulversette 0, Frtisch GmbH, Idar-Oberstein, Germany). Green and senescent leaf samples were stored in scintillation vials at room temperature and analyzed separately. Leaf TC and TN content were determined with a Carlo-Erba NA-1500 elemental analyzer (Fisons Instruments Inc., Danvers, MA, USA). TP was extracted using an acid-digest (HCl) extraction, and concentrations of soluble reactive P were determined by colorimetric analysis (Methods 365.4 and 365.2, US EPA 1983). Leaf C and N bulk isotopic signatures (δ^13^C, δ^15^N) were analyzed on a Thermo Scientific Delta V Plus CF-IRMS coupled to a Carlo-Erba 1108 elemental analyzer via a ConFlo IV interface (Thermo Fisher Scientific, Waltham, MA, USA). All C and N analyses were conducted at the Southeast Environmental Research Center Analysis Laboratory.

Using leaf carbon isotope fractionation values, we calculated the concentration of intracellular CO_2_ and plant water use efficiency integrated over the lifespan of the leaf tissue samples (i.e., intrinsic water use efficiency, *WUE*) via methods described by O’Leary (1988) and Marshall et al. (2007) (and outlined in Lambers et al. 2008). We used an ambient concentration of atmospheric CO_2_ of 408 µmol mol^-1^ for our calculations, which is a conservative estimate for the 2019 calendar year and indicative of the site’s atmospheric conditions based on IRGA measurements from an eddy covariance tower at the site. Thus, the equation used to calculate *c_i_* and *WUE* from carbon isotope data were: *c_i_* = ((−8.5 - δ^13^C - 4.4) ÷ 22.6) × 408), and *WUE* = (408 × (1 - *c_i_* ÷ 408)) ÷ 1.6, where *c_i_* is the value derived from the previous equation. Additionally, the following equation was used to calculate the resorption of N and P using green (G) and senescent (S) leaf nutrient content: Relative resorption (%) = ((G - S) ÷ G × 100) (Pugnaire and Chapin 1993).

### Statistical analyses

Repeated measures analysis of variance (ANOVA) was used to test for differences in water level, surface water salinity, and porewater salinity among locations (fringe and interior), island habitats (center and edge), and season (wet and dry), as well as for the interaction between these effects and season, which was used as the repeated measure. For the repeated measures ANOVA, islands were nested within locations and treated as experimental units. All effects were considered fixed, except for when testing for significant differences in habitat, which included location as a random effect to account for the nested structure of the sampling scheme. One-way ANOVAs were used to test for differences in soil surface elevation among locations and habitats and their interaction. Two-way ANOVAs were carried out for all leaf functional traits and nutrient concentrations, making comparisons across all habitat and season combinations. Tukey HSD posthoc tests were used to identify significant pairwise comparisons when ANOVAs indicated statistical differences. Repeated measures ANOVAs were performed using PROC MIXED (SAS Institute, Cary, NC, USA), and the one-way and two-way ANOVAs were performed in R v3.5.1 (R Core Team 2018).

We constructed linear mixed-effects models (with a Gaussian error distribution and identity link function) to address our research questions. Island habitat and season were included as fixed effects in the models to address questions 1 and 2, respectively, with water levels and porewater salinity being also included as the continuous covariates to parse out their marginal effects. We couple inference from these models to leaf nutrient analyses and our measurements of the hydrological environment to inform about nutrient and water use of *R. mangle* (question 3). Before model fitting, response variables were confirmed to meet the assumptions of data normality. Four separate models were constructed for each of four gas exchange variables of interest: *A_net_*, *g_sw_*, *c_i_*, and *wue*. For each model, fixed effects for season (wet and dry), habitat (center and edge), porewater salinity, and water level were considered, including interaction terms for water level and porewater salinity with season. All models considered random intercept terms for location (i.e., fringe vs. interior), islands, and islands nested within location. Random slopes were explored but determined not to improve model fits. The best-fit models were determined via stepwise model comparison using AIC based on backward selecting random effects then backward selecting fixed effects, as implemented with the ‘lmerStep’ function in the lmerTest R package (Kuznetsova et al. 2017). The best fit models included a random intercept term for islands, which helped remove variability in the data because of the sampling design. Random effects for location were insignificant, signifying that most of the random variance in the gas exchange data was among islands, which we consider as the experimental unit in all mixed-effects models. The mixed-effect models were fit using restricted maximum likelihood estimates via the lme4 R package (Bates et al. 2015). Models were evaluated using model predicting, tabling, and plotting functions from the sjPlot R package (Lüdecke 2018). All analyses were complete in R v3.5.1 (R Core Team 2018).

## RESULTS

### Mangrove-island micro-elevational differences and ecohydrology

Soil surface elevation (measured in relation to the NAVD88 datum) significantly declined from mangrove-island center to edge habitats from −0.14 ± 0.1 m at island centers to −0.4 ± 0.02 m at island edges, a mean difference of about 30 cm (*F*_1,20_ = 108.42, *p* < .001; Table 1). Water levels relative to the soil surface were significantly higher in edge than in center habitats (*F*_1,178_ = 178.33, *p* < .001), measuring on average 36.9 ± 1.4 cm in edge habitats, and 12.8 ± 1.2 cm in mangrove-island centers (Table 2). We recorded water levels of 0 cm (i.e., non-inundated habitats) in 10% of our measurements, and those were exclusive to mangrove-island centers during the dry season (Figure S1C). There was a significant effect of season (*F*_1,178_ = 11.11, *p* < .001) on water levels, where they increased from 17.05 ± 1.5 cm in the dry season to 30.4 ± 1.6 cm in the wet season (Table 2, Figure S1C).

**Table 2.** Seasonal variation in water levels, surface water and porewater salinity measured in mangrove-island habitats at scrub R. mangle dominated mangroves at TS/Ph-7 in southeastern Florida Everglades. Means (± 1 SE) with different letters within each column denoting significant differences among groups (Tukey HSD post hoc, p < .05).

| Season | Habitat | Water level<br>(cm) | Surface water<br>salinity (ppt) | Porewater<br>salinity (ppt) |
| --- | --- | --- | --- | --- |
| Dry | Edge | $33.5 \pm 1.9^A$ | $16.11 \pm 1.16^A$ | $24.22 \pm 0.44^A$ |
| | Center | $10.1 \pm 1.9^B$ | $21.14 \pm 1.51^B$ | $20.83 \pm 0.44^B$ |
| Wet | Edge | $40.2 \pm 1.9^C$ | $15.31 \pm 1.16^A$ | $26.00 \pm 0.44^C$ |
| | Center | $15.5 \pm 1.9^B$ | $14.41 \pm 1.47^A$ | $22.22 \pm 0.44^D$ |

Continuous water level data recorded at the fringe and interior mangrove zones indicated similar flooding trends between locations, with lower water levels during the dry season and higher water levels in the wet season, up to 40-47 cm above the soil surface in both locations (Figure S2). Water levels at the interior mangrove forest always remained higher than those registered in the fringe mangrove zone (Figure S2A). Porewater salinity was significantly different between habitats (*F*_1,178_ = 91.45, *p* < .001) and seasons (*F*_1,178_ = 17.87, *p* < .001), with lower salinity values in the center (21.5 ± 0.3) of the islands relative to the edge (25.1 ± 0.3) habitats, and slightly lower porewater salinity during the dry season (22.5 ± 0.4) than in the wet season (24.1 ± 0.3; Table 2, Figure S1D). There was no significant interaction (*F*_1,178_ = 0.26, *p* > .05) between island habitats and seasons, indicating that the variation in porewater salinity between habitats was independent of seasonality (Table 2). Surface water salinity was not significantly different among center and edge habitats (*F*_1,163_ = 2.36, *p* > .05), but increased significantly from the dry to the wet season (*F*_1,163_ = 8.97, *p* < .01, Table 2). There was also a Tukey posthoc HSD test indicated that only island center habitats in the dry season differed from all other pairwise comparisons (Table 2).

### Rates of leaf gas exchange and their relationships to the hydrological environment

*A_net_* measurements ranged from 0.1 to 15.1 µmol m^-2^ s^-1^, with 90% of the observations recorded between 2 and 14 µmol m^-2^ s^-1^ (see Figure S3). *g_sw_* values were low, ranging from <0.01 to 0.27 and averaging 0.1 mol m^-2^ s^-1^(see Figure S3). Associated *c_i_* values ranged from 40 to 377 and averaged 242 µmol mol^-1^, with 98% of them being greater than 150 µmol mol^-1^. Lastly, measured rates of *wue* varied between >0.01 and 0.21 mmol CO_2_ mol H_2_O^-1^, being normally-distributed about a mean value of 0.09 mmol mol^-1^.

The linear mixed-effects model for *A_net_* included fixed effects for island habitat, porewater salinity, water level, season, and an interaction term for water level with season (Figure S4 & Table S4). There was substantial variation in *A_net_* rates among leaves (σ^2^ of about 6 µmol m^-2^ s^-1^), and the random variation among islands was about 0.02 µmol m^-2^ s^-1^ (see Table S4). All fixed effects were statistically significant (*p* < .05), except the interaction term, which was marginally significant (*p* = .05) but greatly improved model fit. Mangrove edge habitats reduced *A_net_* by over 2.5 µmol m^-2^ s^-1^ relative to mangrove-island centers (Figure 2). Seasonality had a comparable negative effect, leading to an average decrease in *A_net_* of just over 2 µmol m^-2^ s^-1^ in the wet season relative to the dry season (Figure 2). After accounting for variation in the data because of habitat and season, the marginal effects of water level and porewater salinity were positive, albeit weak, leading to increases in *A_net_* of roughly 0.1 µmol m^-2^ s^-1^ per cm increase in water level (Figure 3) or per ppt increase in porewater salinity (Figure 4). Therefore, *A_net_* increased as water levels increased, with increases consistent across habitats (Figure 3); a similar pattern was observed concerning soil porewater salinity, although the magnitude of increase in *A_net_* was smaller (Figure 4). These relationships of *A_net_* with water level variability were consistent across seasons, although rates of *A_net_* were depressed during the wet season (Figure 2). The mixed-effects model for *A_net_* fit satisfactorily for these types of linear mixed-effects models modeling leaf-gas exchange data using environmental predictors, explaining 24% of the variation in the data, 22% of which was explained by ecohydrological data (i.e., fixed effects) (Table S4).

**Figure 2.**
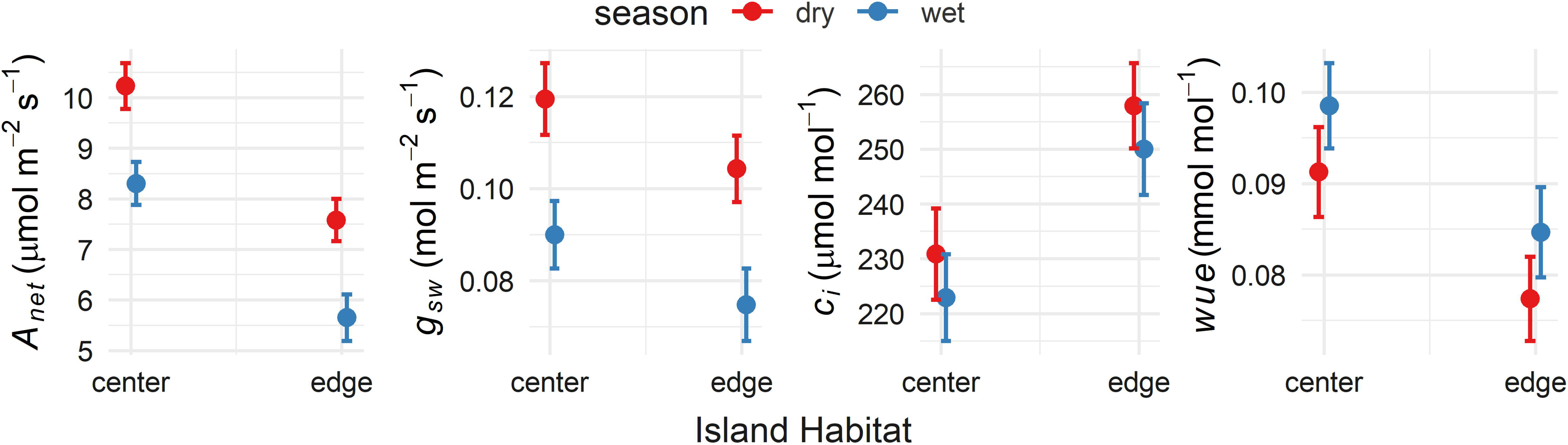
Predicted marginal mean (±95% confidence intervals) values of photosynthesis (*A_net_*), stomatal conductance (*g_sw_*), the concentration of intracellular CO_2_ (*c_i_*), and intrinsic water use efficiency (*wue*) by mangrove-island habitat and season. The dry season is from November to April, and the wet season is from May to October. See supplemental material for complete model summaries.

**Figure 3.**
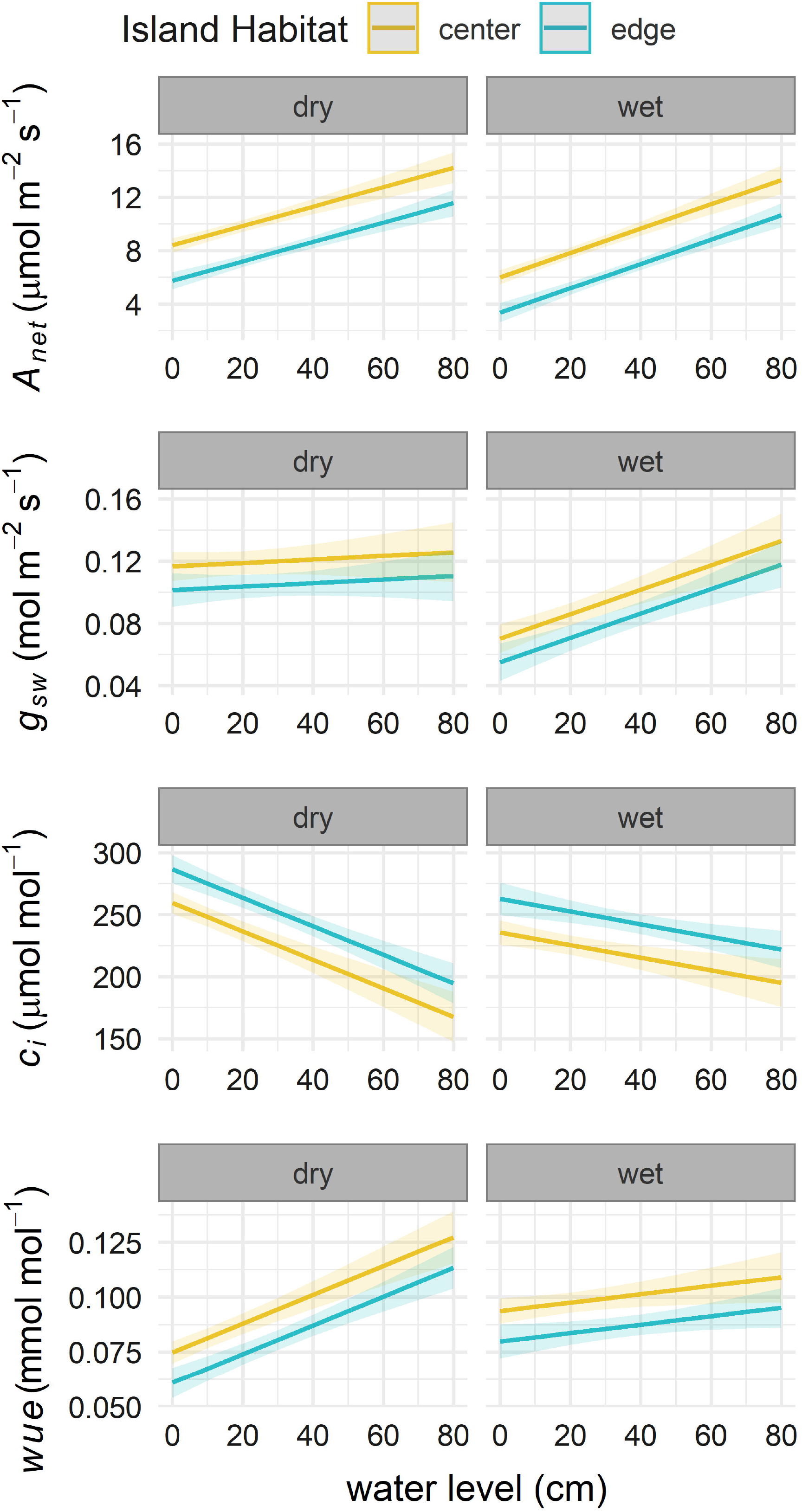
The effect of water level on leaf photosynthesis (*A_net_*) and stomatal conductance (*g_sw_*), the concentration of intracellular CO_2_ (*c_i_*), and intrinsic water use efficiency (*wue*) by season. Lines are habitat-specific predicted mean marginal mean values (± 95% confidence intervals) from linear mixed-effects models.

**Figure 4.**
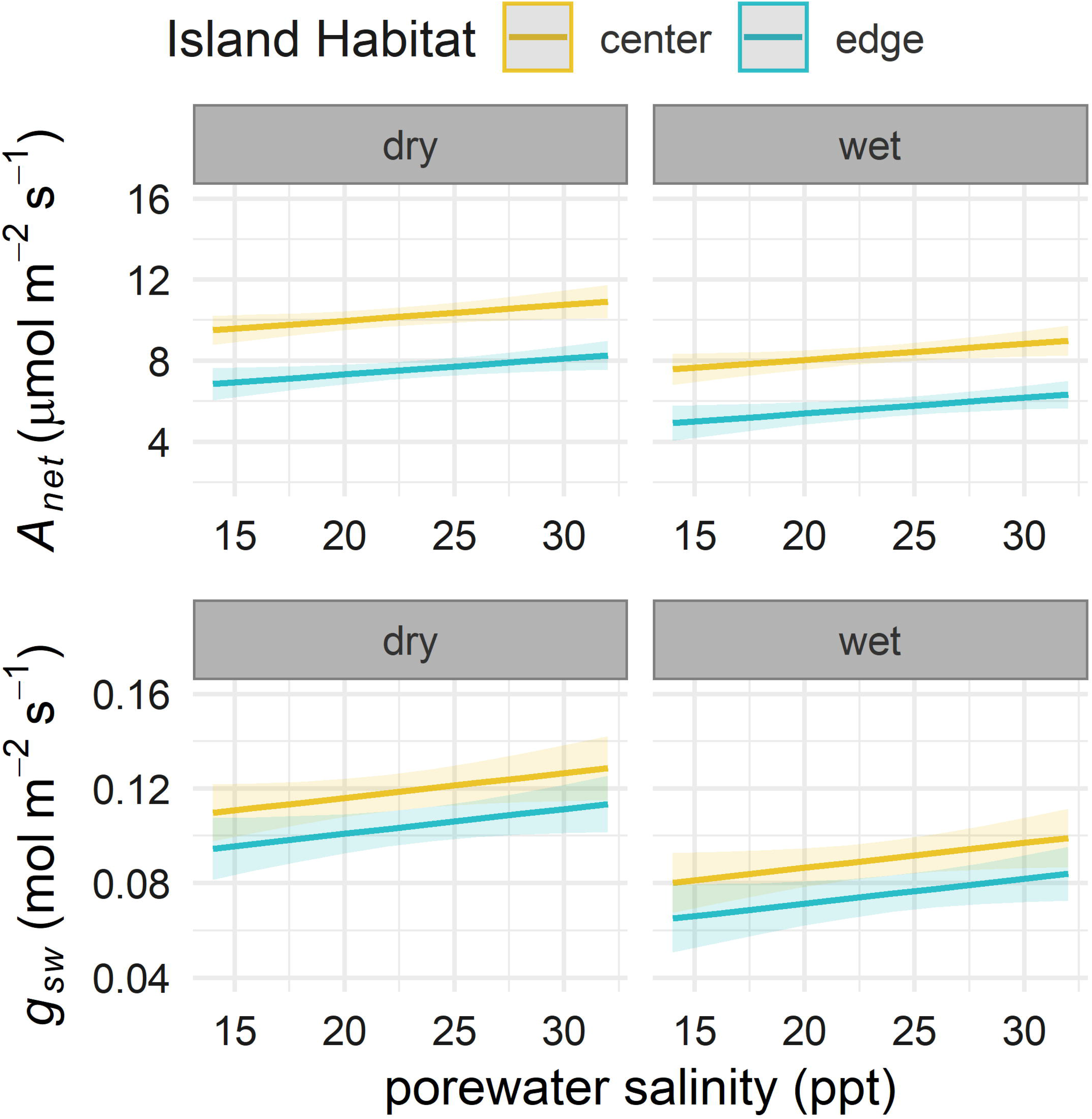
The effect of soil porewater salinity on leaf photosynthesis (*A_net_*) and stomatal conductance (*g_sw_*) by season. Porewater salinity was not included in the best-fit models for *c_i_* or *wue*. Lines are predicted mean marginal effects from linear mixed-effects models ± 95% confidence intervals (colored by island habitat).

*g_sw_* was modeled using an identical mixed-effects model as was used for *A_net_* (Figure S5 & Table S5). Generally, rates of *g_sw_* were low, with 98% of g_sw_ measurements being <0.2 mol m^-2^ s^-1^. Random variance in *g_sw_* among islands was negligible, being <0.01 mol m^-2^ s^-1^. Leaf *g_sw_* in edge habitats was statistically lower than that of mangrove-island centers (*p* < .001), being depressed by about 0.02 mol m^-2^ s^-1^ (Figure 2). Water levels did not affect rates of *g_sw_* (*p* > .05, Figure 3, Table S5), and soil porewater salinity had a marginal effect (*p* = .07) on *g_sw_*, where conductance increased slightly at high salinities, after accounting for the effects of other environmental variables in the model (Figure 4). The effect of season on rates of *g_sw_* was significant in the model, with the wet season leading to a 0.05 mol m^-2^ s^-1^ decrease in conductance (Figure 2) and the interaction between water levels and season being statistically significant (Figure 3). Overall, the mixed-effects model for *g_sw_* did not fit the data as well as the model for *A_net_*. The model only explained about 12% of the variability in the data, with 9% of its explanatory power coming from the environmental predictors (Table S5).

Although the model selection approach was the same as the other mixed-effects models, the best-fit model for *c_i_* differed from the models for *A_net_* and *g_sw_*. The model did not include a fixed effect for soil porewater salinity (which dropped out of the model in the model selection procedure) but included all the same fixed effects as the models for *A_net_* and *g_sw_*, which were all statistically significant (*p* < .001), and a random intercept term for islands (Table S6). Mangrove-island edge habitats had consistently higher *c_i_* values than island centers, being about 27 µmol mol^-1^ greater (19 to 35 µmol mol^-1^ difference in 95% CI estimates, Figure 2). The marginal effect of season alone was similar in magnitude to that of habitat; the wet season led to a decrease in *c_i_* of 24 µmol mol^-1^ (14 to 34 µmol mol^-1^ difference 95% CI estimates) relative to the dry season (Figure 2, Table S6). Water levels, by themselves (again, the marginal effect), led to a slight decrease in *c_i_* but had a positive interaction with season, indicating that the relative decrease in *c_i_* due to increasing water levels was suppressed during the wet season (Figure 3). The random intercept term in the model (for islands) explained a considerable amount of variation in the data (σ^2^ = 128 µmol mol^-1^, with *τ_island_* = 66 µmol mol^-1^). The mixed-effect model for *c_i_* fit the poorest of all four models, explaining just under 12% of the variance in *c_i_*, about 9% of which was explained by data from the hydrological environment (Table S6).

Lastly, we modeled *wue* using a similar mixed-effects model to that of *g_sw_*. In the model for *wue*, all fixed effects were statistically significant (*p* < .001); however, the fixed effects were more-subtle in magnitude. Similar to the model for *c_i_*, porewater salinity was not included in the best-fit model. *wue* values were normally-distributed about a mean value of 0.09 mmol mol^-1^, with 83% of the data having values between 0.05 and 0.15 mmol mol^-1^. Mangrove-island edge habitats had lower *wue* by 0.01 mmol mol^-1^ than island centers (Figure 2). The marginal effect of water level, although being statistically significant in the model, was negligible; however, the wet season caused an increase in *wue* by 0.02 mmol mol^-1^ relative to the dry season, with the interaction between water level and season being slightly negative (Figure 2, Figure 3). Random variation in *wue* structured across the eight mangrove-islands was minuscule, being <0.01 mmol mol^-1^. Thus, the model fit for *wue* was comparable to, and slightly better than, the model for *c_i_*, with fixed effects explaining just over 12% of the variance in the data, about 9% of which was explained using the environmental predictors (Table S7).

### Rhizophora mangle leaf functional traits, nutrient content, and isotopic signatures

Leaf SLA values did not vary significantly between seasons (*F*_1,155_ = 0.46, *p* > .05) and island habitats (*F*_1,155_ = 3.07, *p* > .05, Table 3), despite having some variation in SLA with average values ranging from 29 to 40 cm^2^ g^-1^. Similarly, leaf water content was not significantly different between all season-habitat combinations (*F*_1,155_ = 0.32, *p* > .05), despite a statistically significant effect of season alone (*F*_1,155_ = 9.10, *p* < .01), where leaf water content was greater in the dry season (65.6 ± 0.3%) relative to the wet season (63.8 ± 0.4%, Table 3).

**Table 3.** Leaf functional traits, carbon and nutrient contents and N:P ratios, nitrogen and phosphorus resorption efficiencies, bulk isotopic signatures, intrinsic intracellular CO_2_ concentrations (*c_i_*) and intrinsic water use-efficiency (*WUE*) (calculated from ^13^C fractionation) for scrub *R. mangle* leaves collected from mangrove-island habitats at TS/Ph-7 during the dry and wet seasons of 2019. Means (± 1 SE), with different letters across each row denoting significantly different groups (Tukey HSD test, *p* < .05).

| Leaf Trait | Dry season |  | Wet season |  |
| --- | --- | --- | --- | --- |
|  | Edge | Center | Edge | Center |
| Leaf dry mass (g) | 0.64 ± 0.02<sup>A</sup> | 0.65 ± 0.02<sup>A</sup> | 0.69 ± 0.02<sup>A</sup> | 0.70 ± 0.02<sup>A</sup> |
| Leaf area (cm<sup>2</sup>) | 24.9 ± 0.7<sup>A</sup> | 25.5 ± 0.7<sup>A</sup> | 26.0 ± 1.4<sup>A</sup> | 27.3 ± 0.9<sup>A</sup> |
| SLA (cm<sup>2</sup> g<sup>-1</sup>) | 28.97 ± 0.66<sup>A</sup> | 39.79 ± 0.68<sup>A</sup> | 37.33 ± 0.40<sup>A</sup> | 40.04 ± 1.71<sup>A</sup> |
| LWC (%) | 65.7 ± 0.4<sup>A</sup> | 65.4 ± 0.5<sup>A</sup> | 63.6 ± 0.5<sup>A</sup> | 64.0 ± 0.8<sup>A</sup> |
| Total C (mg g<sup>-1</sup>) | 450.9 ± 5.7<sup>A</sup> | 441.4 ± 1.0<sup>A</sup> | 428.7 ± 17.7<sup>A</sup> | 444.0 ± 1.0<sup>A</sup> |
| Total N (mg g<sup>-1</sup>) | 9.8 ± 0.2<sup>AB</sup> | 10.2 ± 0.3<sup>A</sup> | 8.4 ± 0.5<sup>B</sup> | 9.0 ± 0.3<sup>AB</sup> |
| Total P (mg g<sup>-1</sup>) | 0.50 ± 0.01<sup>AB</sup> | 0.55 ± 0.04<sup>A</sup> | 0.42 ± 0.01<sup>B</sup> | 0.46 ± 0.02<sup>AB</sup> |
| Leaf N<sub>area</sub> (mg cm<sup>-2</sup>) | 0.258 ± 0.012 | 0.264 ± 0.019 | 0.263 ± 0.035 | 0.306 ± 0.020 |
| Leaf P<sub>area</sub> (mg cm<sup>-2</sup>) | 0.013 ± 0.001 | 0.014 ± 0.002 | 0.013 ± 0.001 | 0.016 ± 0.001 |
| Atomic N:P | 41.7 ± 1.9 | 40.0 ± 0.1 | 43.5 ± 3.7 | 42.5 ± 2.1 |
| N resorption (%) | 60.0 ± 0.4 | 62.8 ± 0.5 | 60.8 ± 2.8 | 62.9 ± 1.5 |
| P resorption (%) | 78.6 ± 0.2 | 74.3 ± 3.2 | 75.5 ± 0.4 | 73.2 ± 6.2 |
| δ<sup>13</sup>C (‰) | -25.5 ± 0.1<sup>AB</sup> | -25.1 ± 0.1<sup>A</sup> | -25.8 ± 0.1<sup>B</sup> | -25.9 ± 0.2<sup>AB</sup> |
| *c<sub>i</sub>* (μmol mol<sup>-1</sup>) | 228.1 ± 2.2<sup>AB</sup> | 219.9 ± 1.8<sup>B</sup> | 233.3 ± 1.1<sup>A</sup> | 235.3 ± 3.6<sup>A</sup> |
| *WUE* (mmol mol<sup>-1</sup>) | 0.1124 ± 0.0014<sup>AB</sup> | 0.1175 ± 0.0012<sup>B</sup> | 0.1092 ± 0.0007<sup>A</sup> | 0.1080 ± 0.0022<sup>A</sup> |
| δ<sup>15</sup>N (‰) | -5.3 ± 0.5<sup>B</sup> | -0.4 ± 0.4<sup>A</sup> | -4.2 ± 1.0<sup>AB</sup> | -0.8 ± 1.5<sup>A</sup> |

Leaf TC content ranged from 400 to 450 mg g^-1^ (Table 3) and was not different between seasons (*F*_1,12_ = 1.10, *p* > .05), habitats (*F*_1,12_ = 0.10, *p* > .05), or their interaction (*F*_1,12_ = 1.77, *p* > .05). Leaf TN concentrations were higher in the dry season compared to the wet season (*F*_1,12_ = 11.95, *p* < .01) and ranged from 8-10 mg g^-1^ (Table 3). There was neither a significant difference in leaf TN between habitats (*F*_1,12_ = 1.86, *p* > .05), nor a significant interaction between seasons and habitats (*F*_1,12_ = 0.11, *p* > .05, Table 3). Leaf TP content did vary significantly between seasons (*F*_1,12_ = 15.05, *p* < .01) and had marginally-significant difference between habitats (*F*_1,12_ = 4.55, *p* = .054), but the interaction effect was not significant (*F*_1,12_ = 0.08, *p* > .05). Overall, mean leaf TP values ranged from 0.42 to 0.55 mg g^-1^ across seasons and habitats, with higher concentrations during the dry season than in the wet season and higher leaf tissue TP values in the island center habitats compared to edge habitats (Table 3, Figure 5). Mean N resorption for *R. mangle* leaves was similar across seasons and habitats and ranged from 60 to 63% (Table 3, Figure 5). In contrast, P resorption of leaf tissue had a broad range compared to that of N, ranging from 73.2 ± 6.2% (center, wet season) to 78.6 ± 0.2% (edge, dry season) across seasons and habitats. Overall, P resorption of *R. mangle* leaves was higher in the edge habitats relative to the center during both seasons (Table 3).

**Figure 5.**
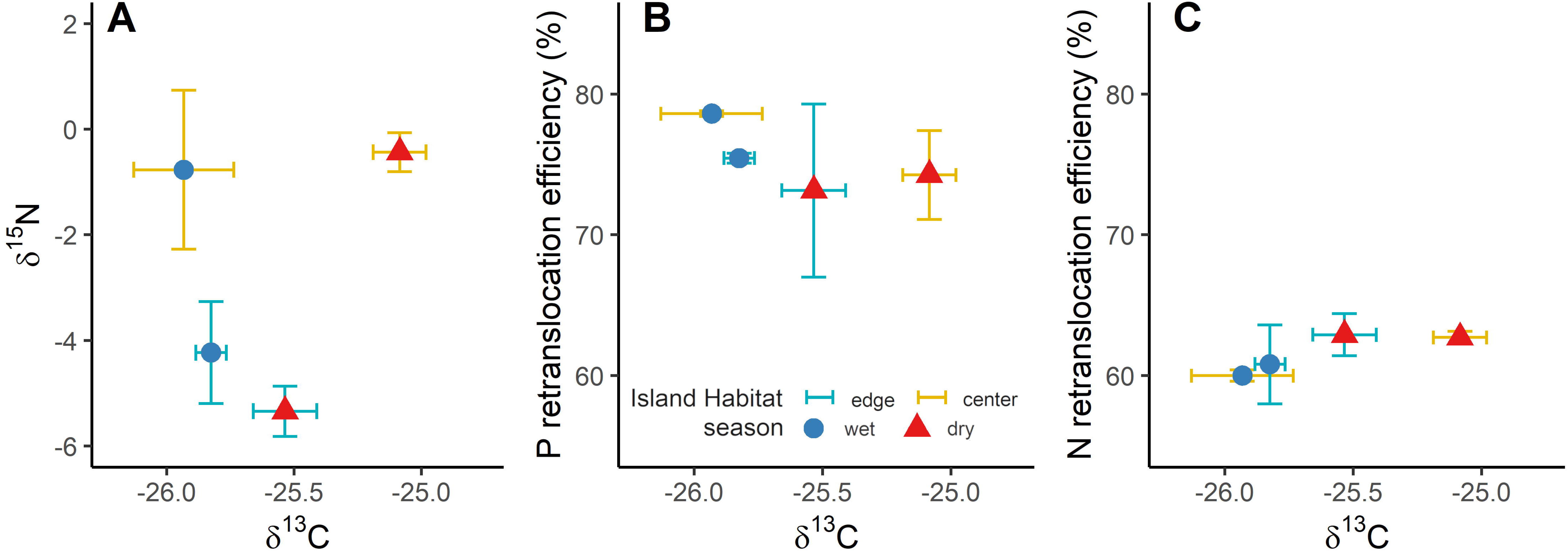
Mean (± 1 SE) leaf isotopic signatures and nutrient resorption efficiency by island habitat and season combination. **A)** the relationship between δ^15^N and δ^13^C in *R. mangle* leaves, **B)** the relationship between N resorption efficiency and δ^13^C for *R. mangle* leaves, and **C)** the relationship between P resorption efficiency and δ^13^C for R. mangle leaves. Error bars colors denote island habitats, while point symbols and colors show seasons.

Patterns in green leaf carbon isotope signatures (δ^13^C) mirrored those of leaf TN and TP concentrations. Carbon isotopic fractionation was more negative during the wet season than in the dry season (*F*_1,12_ = 18.88, *p* < .01, Table 3, Figure 5), with no statistical difference between habitats (*F*_1,12_ = 1.17, *p* > .05). Green leaves bulk δ^13^C values ranged from −25.9 to −25.1‰ across seasons and habitats (Table 3, Figure 5). Physiologically, the differences in carbon isotopic fractionation were estimated to result in a maximum difference of about 10 µmol mol^-1^ *c_i_* between the center and edge habitats and a difference of 5 to 15 µmol mol^-1^ *c_i_* within habitats (*F*_1,12_ = 1.71, *p* > .05) because of seasonality (*F*_1,12_ = 18.88, *p* < .01). These differences resulted in slightly greater, but not statistically different, *c_i_* values in mangrove-island centers than in edge habitats in the wet season; however, the opposite pattern was found during the dry season, with *c_i_* being about 10 µmol mol^-1^ greater in island edge habitats relative to their centers (Table 3). The interaction between season and habitat was marginally significant (*F*_1,12_ = 4.53, *p* = .055). Intrinsic water-use efficiency (*WUE*) was calculated from leaf δ^13^C values; accordingly, *WUE* was greatest in mangrove-island centers during the dry season relative to all habitat-season combinations. Additionally, *WUE* was significantly lower in the wet season than the dry season (*F*_1,12_ = 18.88, *p* < .01, Table 3). Mean leaf bulk δ^15^N values were significantly higher (*F*_1,12_ = 19.66, *p* < .001) in the center habitats (−0.60 ± 0.66‰) relative to the edge (−4.79 ± 0.66‰), but there was no difference between seasons (*F*_1,12_ = 0.17, *p* > .05) and no interaction between season and island habitat (*F*_1,12_ = 0.58, *p* > .05, Table 3, Figure 5A).

## DISCUSSION

Our first research question asked how mangrove-island habitat affects rates of leaf gas exchange. We can confirm our hypothesis that photosynthetic rates and stomatal conductances are greater at island centers than edges (Figure 2). However, contrary to our expectation, habitat driven-variation in leaf gas exchange rates was roughly equal to seasonal variation, with no apparent decoupling between *A_net_* and *g_sw_* (Figure 2). Our second research question asked whether water levels or salinity exerted a more-substantial effect on mangrove leaf gas exchange *in situ*. Porewater salinities at TS/Ph-7 are relatively low (i.e., between 15 and 30 ppt), compared to the levels of salinity at which leaf gas exchange rates of *R. mangle* are negatively affected (i.e., salinities > 35 ppt) and did not vary considerably over time (Figures S1 & S2). Therefore, we conclude that inundation stress is the primary driver of variation in *R. mangle* leaf gas exchange rates. Lastly, we predicted that general physiological stress would be lower at island centers than island edge habitats, leading to increased *wue* and higher rates of nutrient resorption at centers relative to edges. Indeed, intrinsic water-use efficiency was greater at island centers than edge habitats, with results being consistent across gas exchange-measured *wue* and isotope-derived *WUE.* In addition, *w*ater levels modulated leaf intrinsic water use efficiency (Figure 3, Table 2), especially in the dry season. Patterns of nutrient resorption were less clear but seemed to indicate differences in leaf N and P concentrations on mangrove-island centers versus edges, illustrating differences in water and nutrient use of *R. mangle* among habitats, which likely drive variation in leaf gas exchange rates.

### The effect of mangrove-island habitat on leaf gas exchange

Our results showed significant differences in soil elevation of about 30 cm between mangrove-island habitats (Table 1), which affected *R. mangle* leaf gas exchange rates (Figures 2, 3, 4). The soil elevation gradient at our study site is driven by differences in mangrove root biomass and productivity between center and edge island habitats, with higher total root biomass and productivity (top 0-90 cm of soil) observed in center habitats compared to the edge habitats (Castañeda-Moya et al. 2011). Along this micro-elevation gradient, we measured clear differences in *A_net_* and *g_sw_* (Figure 2). *A_net_* was nearly 3 µmol m^-2^ s^-1^ (or 20%) greater at mangrove-island centers than edges, and *g_sw_* was >0.1 mol m^-2^ s^-1^ (or >37%) higher; these differences were attributable to mangrove-island habitat alone, after accounting for variation explained by water level, salinity, or seasonality (i.e., marginal differences). Associated *c_i_* concentrations were about 30 µmol mol^-1^ (or 12%) lower, and *wue* was >0.01 mmol mol^-1^ (or about 10%) greater at island centers than at island edges (Figure 3).

Thus, these findings support our first hypothesis about the effect of habitat micro-elevation (center vs. edge) on *A_net_*, with overall greater leaf gas exchange rates at mangrove-island centers compared to their edges. Interestingly, the effect of habitat on *R. mangle* leaf gas exchange rates was similar in magnitude to the effect of season (Figure 3). The magnitude of variation in *A_net_* that we report in this study is slightly larger than the magnitude of variation reported by Lin and Sternberg (1992), who found that *A_net_* varied up to 2 µmol m^-2^ s^-1^ between scrub and fringe *R. mangle* trees in the nearby Florida Keys. Furthermore, our *A_net_* measurements with average values between 5.7 µmol m^-2^ s^-1^ (edge habitat, wet season) and 10.2 µmol m^-2^ s^-1^ (center habitat, dry season, Figure 2), are within the range of values reported for *R. mangle* interior scrub (5.3 µmol m^-2^ s^-1^) and fringe (10 µmol m^-2^ s^-1^) mangroves along a distinct zonation pattern in the intertidal zone at Twin Cays, Belize (Cheeseman and Lovelock 2004). Island center habitats may also support greater access to mixed soil-groundwater sources in the dry season facilitating higher leaf gas exchange rates because of increased freshwater availability, or a reduction in the energy demand for processing saline water (Ewe et al. 2007). Our results demonstrate the effect that higher-elevation center habitats at TS/Ph-7 have on alleviating inundation stress, which pervades scrub mangrove physiology, making trees growing in center habitats in the dry season physiologically comparable to fringe mangroves. Certainly, the stress relief is short-lived when water levels rise in the wet season (Table 2, Figure S1), and leaf gas exchange rates are depressed once more (Figure 2. Figure S3).

### Seasonal signals in R. mangle physiology with implications for ecosystem functioning

We found that *A_net_* varied over 2.5 µmol m^-2^ s^-1^ (17%), and *g_sw_* varied about 0.03 mol m^-2^ s^-1^ (11%) within habitats between the wet and dry seasons (Figure 3). Despite differences in *A_net_* and *g_sw_* between seasons, we found no statistical differences in *c_i_* and *wue* between seasons, although there was some variation (Figure 3). These differences point to habitat-specific optimization of the diffusion and uptake of CO_2_ into (i.e., *c_i_*) and the movement of water vapor out of (i.e., *wue*) leaves (Cardona-Olarte et al. 2006, Barr et al. 2009, Reef and Lovelock 2015, Lopes et al. 2019). As precipitation and freshwater flow increased during the wet season, water levels increased, and mangrove-island centers experienced greater inundation levels (Figures S1 & S2), resulting in decreased *A_net_* and *g_sw_* (Figure 3). A similar reduction in *A_net_* and *g_sw_* was measured in mangrove-island edge habitats during the wet season (Figures 3 & 4). Although *A_net_* was depressed in the wet season, the effect of inundation levels on reducing *A_net_* was consistent across seasons (Figure 4). *g_sw_* showed a similar pattern to *A_net_*, being highest in mangrove-island centers during the dry season (Figure 3). However, the effect of water levels on *g_sw_* resulted in increased *g_sw_* in the wet season, an effect which was tempered during the dry season (Figure 4).

In the Florida Everglades, irradiance peaks in April and May (Barr et al. 2009), and rainfall and temperature reach maxima in June, July, and August (Figure 1 A, B). Thus, photosynthetic demand for water is likely highest from April to May, at the end of the dry season and the beginning of the wet season. During this time, we measured lower water levels and porewater salinities relative to the peak wet season. Barr et al. (2009) recorded earlier diurnal and more considerable reductions in *g_sw_* during late May versus July or August for mangroves at Key Largo, evidencing the greatest water-limitation on photosynthesis occurs at the end of the dry season. Additionally, the greatest *A_net_* rates for tall fringe mangroves in the southeastern Everglades occur during the dry season from March to May (Barr et al. 2009). The difference in surface water and porewater salinity (*Δsw-pw*) can be used as a proxy for tree transpiration (Reef and Lovelock 2015). Average *Δsw-pw* measured 10.7 and 8.1 ppt at mangrove-island edges in the wet and dry seasons, respectively, whereas it measured 7.8 and −0.3 in mangrove-island centers in the wet and dry seasons, respectively. Indeed, measured transpiration was highest at the end of the dry season in March and April (Figure S3). Thus, photosynthetic demand for water is higher in the dry season in mangrove-island centers relative to edges or either habitat in the wet season. The drying of the soils at slightly higher-elevation island center habitats in this scrub mangrove forest likely facilitates increases in *A_net_*. Therefore, the seasonal variation in hydrology, mainly reductions in water levels and porewater salinity during the dry season, albeit coupled with an increase in surface water salinity in this study (Table 2), likely have critical consequences for mangrove forest carbon fluxes at greater spatial scales. Potentially drying soils could also lead to an increase in ecosystem respiration (Chambers et al. 2014), or non-stomatal derived CO_2_ uptake (Reef and Lovelock 2015). Future research could look at soil metabolic dynamics (e.g., soil respiration, microbial C and N, or changes in microbial communities) with hydrology and season, which may show unique responses in this scrub *R. mangle* forest (Lovelock 2008, Chambers et al. 2014).

### The effect of salinity and water level on R. mangle leaf gas exchange

During 2019, the hydrological environment (Figure 1C, D) at our study site was seasonally dynamic (Table 2) and tended to mirror patterns in local rainfall (Figure 1B). Water levels and porewater salinity both increased during the wet season (Table 2, Figure S1) from the beginning of the rainy season in May through November. This likely led to increased water column stratification via a larger freshwater lens (Hughes et al. 1998, Uncles et al. 1992). Indeed, the difference in surface water and porewater salinity increased in the wet season, with surface water salinities decreasing, despite a slight increase in porewater salinities (Table 2, Figure S1). When data were grouped by season, edge habitats were slightly more saline (about 4 ppt on average) than mangrove centers (Table 2), and there were no apparent differences between fringe and interior scrub mangrove zones (Figures S1 & S2). Comparing these changes in the hydrological environment with previous years, long-term water level and porewater salinity data at this site show that water level usually increases and porewater salinity usually decreases in the wet season relative to the dry season (Castañeda-Moya et al. 2013). We measured the opposite trend for porewater salinity in 2019 with slight differences between seasons, likely because it was a wet year. Total rainfall for 2019 (929 mm yr^-1^; Figure S1) was 10% greater than the total for rainfall 2018 (859 mm yr^-1^; https://sofia.usgs.gov/eden).

Rates of mangrove leaf gas exchange (i.e., *A_net_* and *g_sw_*) typically decrease with porewater salinity, especially along strong salinity gradients in the environment (i.e., gradients >30 ppt, Ball 2009, Clough and Sim 1989, Lugo et al. 2007). Porewater salinity was only included in the linear mixed-effects models for *A_net_* and *g_sw_*, and its marginal effect was minimal, slightly increasing *A_net_* by about 0.1 µmol m^-2^ s^-1^ per ppt increase in porewater salinity. Like the effect of porewater salinity on *A_net_*, the effect of porewater salinity on *g_sw_* was small in magnitude and consistent across seasons and mangrove-island habitats but was not statistically significant (Figure 4). The minimal influence of porewater salinity on leaf gas exchange is likely due to the minor seasonal and spatial variation in salinity that we measured during 2019. Differences were not large, maximizing at 16.4 ppt and averaging 5.2 ppt, especially when considering that *R. mangle* frequently occupies natural habitats with salinities greater than seawater (Reef and Lovelock 2015), potentially up to 50-60 ppt (Cintron et al. 1978). At our study site, variation in porewater salinity from long-term monitoring data (2001-2020) has shown similar magnitudes of relatively-minor variation in porewater salinity, with overall mean values ranging from 19-22 and rarely exceeding 30 ppt (Castañeda-Moya et al. 2013). Additionally, long-term variation in porewater salinity (<30 ppt) across the FCE mangrove sites (Shark and Taylor River sites) is below the critical value of 65 ppt that influences forest structure and productivity across the FCE landscape (Castañeda-Moya et al. 2013). Thus, the limited effect of salinity in our linear mixed-effects models is likely broadly indicative of relatively weak salinity effects in both scrub and tall *R. mangle*-dominated forests of the Everglades. This is a significant finding, given that these scrub forests are adapted to relatively low salinities. If salinity increases greatly due to SLR and saltwater intrusion in the region, they will likely experience more stressful conditions with could diminish their physiological performance, as observed in other studies in the neotropics (Lugo et al. 2007).

The effects of inundation on *R. mangle* photosynthesis can be difficult to separate from the effects of salinity; however, the linear mixed modeling approach we used permitted doing so. We found that the intermittent flooding conditions of mangrove-island centers that averaged 10-15 cm above the soil surface allowed greater *A_net_* and *g_sw_* than permanently flooded mangrove-island edges, which averaged 30-40 cm water levels. This indicates that the hydrological regime in center habitats allows mangrove soils to repeatedly flood and desiccate, which may help the species maintain optimal stem water potentials and *g_sw_* (Ball 2009, Reef and Lovelock 2015). In typical greenhouse experiments where mangrove seedlings are grown, inundation alone has little effect on photosynthetic rates or biomass production (Pezeshki et al. 1990b, Hoppe-Speer et al. 2011). However, inundation may sometimes lead to increases in leaf gas exchange rates over the short term and often interacts with salinity over time to reduce *A_net_*, *g_sw,_* and growth rates (e.g., Cardona-Olarte et al. 2013). Thus, water levels and flooding duration are key drivers controlling *A_net_* in mangroves, and mangroves seem to physiologically\ optimize photosynthesis to water levels. For instance, findings from a long-term greenhouse inundation study by Farnsworth and Ellison (1996) exemplify how short-term responses of *R. mangle* to inundation differ from longer-term responses. Over several years, high inundation levels led to steady declines in *A_net_* of up 25% for a given *g_sw_* and decreases in growth rates. Results of the high-water level (30-40 cm above soil surface) treatment were similar to those of the low water level (10-15 cm) treatment, suggesting that *R. mangle* physiology is optimized at inundation levels that reach just a few centimeters above the soil surface at high water level (Ellison and Farnsworth 1997). Further research could assess the coupled effect of the depth and duration of flooding with salinity in the FCE by characterizing photosynthetic rates across a landscape-scale gradient that encompassed multiple sites and a broad range of salinity (or potentially with an experimental increase of salinity).

Although initial increases in *R. mangle g_sw_* can result from short term inundation (Krauss et al. 2006, Hoope-Speer et al. 2011), especially at low salinities (Pezeshki et al. 1990b), several studies have linked stomatal closure to longer-term inundation (Kozlowski 1997, Ellison and Farnsworth 1997). We measured depressed *g_sw_* during the wet season and in mangrove-island edge habitats relative to centers; however, this was not attributable to water levels after accounting for variation in seasonality and habitat, in that *g_sw_* increased with increasing water levels during the wet season. Our measurements of *g_sw_* were consistent with those reported in other studies from across a range of inundation levels (Clough and Sim 1989, Lin and Sternberg 1992, Ellison and Farnsworth 1997, Krauss et al. 2006, Lugo et al. 2007, Barr et al. 2009), supporting the understanding that *R. mangle* leaves limit *g_sw_* in response to flooding. Limits on *g_sw_* seek to optimize *c_i_* for carbon gain without losing unnecessary amounts of water, but our findings show that *g_sw_* can increase with freshwater inputs, resulting in a decrease in *c_i_* as *A_net_* increases (Figure 3), likely because of faster Calvin cycle reactions (see supplemental information Figure S9). Interestingly, the scrub *R. mangle* leaves of TS/Ph-7 operate with low *c_i_* concentrations (range = 220-260 µmol mol^-1^), which suggests pervasive inundation stress. Such pervasive inundation stress likely leads to water and nutrient (i.e., leaf N and Rubisco)-stressed photosynthesis, which decreases max *A_net_* at mangrove-island edges by reducing maximum rates of carboxylation, especially in the wet season.

### R. mangle nutrient use at TS/Ph-7

We found little variation (60-63%) in N resorption efficiencies for *R. mangle* leaves across scrub mangrove-island habitats (Figure 5). In contrast, higher overall efficiencies of P resorption (73-79%) of leaf tissue were measured across habitats, with higher P resorption in mangrove-island edge habitats relative to centers, suggesting higher P availability in island centers. Our findings are roughly comparable to N and P resorption efficiencies for *R. mangle* in the control plots of scrub-dominated forests in Panama (∼50%, and 80%, respectively; Lovelock et al. 2004). Resorption of nutrients from senescent leaves before leaf fall is a within-stand nutrient recycling mechanism that may reduce nutrient losses via tidal export in coastal systems (Vitousek 1982, Aerts and Chapin 2002). Like other tropical trees, mangroves exhibit several physiological mechanisms that reduce nutrient losses via tidal exchange, including resorption of N and P before leaf abscission, which can lead to increased availability of limiting nutrients and ultimately change nutrient use and conservation patterns (Twilley et al. 1986, Alongi et al. 1992, Feller et al. 2003a, 2003b).

Our findings suggest that *R. mangle* conserves P better than N in this P-limited environment and indicate that canopy N and P resorption efficiency at TS/Ph-7 potentially results from the differential acquisition of these nutrients from the soil between habitats, and variation in the use of these nutrients among leaf stages. Mangrove species prioritize resorption of nutrients that are limited in the soil, and it has been suggested that plants growing in nutrient-poor environments resorb a higher proportion of nutrients, potentially decreasing nutrient loss by efficient nutrient recycling (Chapin and Moilanen 1991). At our study site, low soil TP concentrations probably determine the higher recycling efficiency of P relative to N. Indeed, soil TP concentrations in the upper 50 cm of soil at TS/Ph-7 (0.06 ± 0.004 mg cm^-3^) are three times lower than soils at the mouth of Shark River estuary (SRS-6), which is dominated by fertile well-developed tall riverine mangroves. Such low TP concentrations result in extreme P limitation at TS/Ph-7 with average soil N:P ratios of 102 ± 6 (Castañeda-Moya et al. 2013). Therefore, the heterogeneous distribution of essential nutrients within mangrove habitats creates distinct nutrient gradients and hot spots along the intertidal zone, influencing the efficiency of internal nutrient recycling. This is supported by observations that nutrient resorption efficiencies in mangroves vary with nutrient availability, e.g., via nutrient addition (Feller et al. 1999, Feller et al. 2003b) or along natural fertility gradients (Medina et al. 2010). Such variation in nutrient availability and resorption efficiencies within mangrove trees likely scales with variation in photosynthesis and productivity (e.g., growth, litterfall) and carbon residence times (e.g., soil and biomass dynamics) of mangrove forests.

Foliar δ^15^N values integrate long-term processes of N sources because isotopic fractionation against the heavier isotope (i.e., ^15^N) occurs during N transformations and interactions between biotic (e.g., mycorrhizal fungi, or bacteria) and biogeochemical (e.g., nitrification, denitrification) nutrient cycling processes (Garten 1993). In our study site, patterns of δ^15^N in *R. mangle* leaves differed drastically between mangrove habitats, with values around −4 to −5‰ for mangrove edge habitats and between 0 and −1‰ for island centers, indicating lower ^15^N discrimination in island center habitats (Table 3, Figure 5A). These δ^15^N values are considerably more depleted than the *R. mangle* leaf δ^15^N values reported for riverine mangroves along Shark River estuary (He et al. 2020), where values were negatively correlated with distance inland from the mouth of the estuary with more enriched leaves occurring near the mouth of Shark River (4‰) relative to upstream (0.4‰) regions. Reported δ^15^N values for *R. mangle* leaves across different ecotypes in the neotropics range from 0 to −11‰, with more negative values for scrub mangrove forests (−5 to −10‰) than for fringe mangroves (0-7‰; Reis et al. 2017a). Similarly, Medina et al. (2010) showed that leaves from interior scrub mangrove communities had more negative δ^15^N values than tall fringe mangroves in eastern Puerto Rico (−12‰ vs. 0‰, respectively). Those δ^15^N values were more negative than those reported for scrub *R. mangle* forests in Florida (Fry et al. 2000), Belize (McKee et al. 2002, Wooller et al. 2003, Fogel et al. 2008), or Brazil (Reis et al. 2017b).

Patterns of foliar δ^15^N between mangrove ecotypes can be discerned using *in situ* leaf nutrient content. For example, a direct relationship between ^15^N discrimination and leaf N:P ratios of *R. mangle* leaves previously reported for the six FCE mangrove sites, including our study site, indicates that leaf N:P ratios accounted for 70% of the variability in ^15^N discrimination (Mancera-Pineda et al. 2009). Thus, foliar ^15^N composition can reflect *in situ* leaf N-status and differences in plant N-use. Hypoxic conditions in the soil may inhibit denitrification and ammonia volatilization, two processes that enrich the soil substrate in ^15^N (Craine et al. 2015). Therefore, the substrate should be less enriched at mangrove-island edges relative to their centers, because of interactions with the soil and the open water channels, which can alleviate hypoxia. Thus, it appears that more negative ^15^N values in the edge habitats may be associated with lower inorganic N (i.e., porewater ammonium) use by edge mangrove trees compared to those in center island habitats (Fry et al. 2000). However, our results contrast slightly with those of Mancera-Pineda et al. (2009), who reported mean δ^15^N values of +3 from 65 mature leaves collected in 2001 at our study site. We posit that differences in δ^15^N values between the two studies could be attributed to the location where leaves were collected during the 2009 study, concluding that it is very likely that Mancera-Pineda et al. only collected leaves from the center of mangrove-islands, avoiding edge habitats. Taking this into consideration, mangrove-island centers potentially may have even more-positive δ^15^N signatures than we found, illustrating that in the center of mangrove-islands, N is taken up by roots in inorganic soluble forms (e.g., porewater ammonium, nitrate) and not biotically via root symbionts.

Another potential explanation of why δ^15^N values were more negative at mangrove-island edges than in their centers is because lateral surface roots of *R. mangle* can extend into open water where they associate with symbiotic biofilms (i.e., algae and aquatic bacteria) that facilitate N acquisition from open water (Potts 1979). A significant source of isotopic discrimination occurs during N transfer between belowground symbionts (e.g., mycorrhizal fungi or bacteria) and plant roots during nitrification, denitrification, and ammonia volatilization. The lighter isotope ^14^N reacts faster than ^15^N (i.e., it is preferentially given to the host plant by the symbiont) so that plant tissues are depleted while substrates are enriched (Högberg 1997, Robinson 2001). Indeed, at our study site, we observed several long, absorptive, fine lateral root systems that protruded from the edge of mangrove-islands into the open water ponds, which were colonized by algal biofilms. Mangrove trees can potentially adapt to nutrient shortage or localized nutrient deficiencies in the soil by altering patterns of nutrient use. This plant strategy may maximize the efficiency of capturing limiting resources essential for growth (e.g., N, P) from soil or surrounding open water areas in nutrient-poor environments such as Taylor River, as proposed by the optimal plant allocation theory (Chapin et al. 1987, Gleeson and Tilman 1992). We observed a slight decrease in foliar δ^15^N during the wet season (Figure 5A, Table 3) as water levels and porewater salinity increased, suggesting that N-acquisition by *R. mangle* via algal biofilms may be slightly greater in the dry season than in the wet season. Lastly, highly depleted (i.e., negative) N-isotope values in leaf tissues are characteristic of tropical wetlands with P limitation because P limitation increases N fractionation, especially in flooded wetlands with limited P pedogenesis (McKee et al. 2002, Troxler 2007, Medina et al. 2008). This is likely the case with the scrub *R. mangle* forest at TS/Ph-7, where the main source of P is brackish groundwater discharge (Price et al. 2006). Soil total P concentrations in the top 10 cm of the peat soils at this site have measured 0.055 (± 0.01) mg cm^-3^, with atomic N:P ratios of roughly 72 (± 2) (Mancera-Pineda et al. 2009), which is considerably lower than soils of most mangrove forests globally, but consistent with mangrove forests in karstic environments (Rovai et al. 2018).

## CONCLUSIONS

Habitat heterogeneity, resulting from micro-elevational differences in mangrove tree locations on islands within the open water, mangrove-island forest landscape, drives variation in scrub *R. mangle* leaf physiological performance. In particular, mangrove-island edge habitats experience greater and more-prolonged inundation than island centers in a seasonal dynamic, which leads to reductions in *g_sw_,* reduced *A_net,_* and slightly lower *wue*. Conversely, mangrove-island center habitats are alleviated from inundation stress in the dry season, leading to increases in *A_net_* and *g_sw_*. Interestingly, *c_i_* levels increase with increasing water levels because inundation likely slows not only *g_sw_*, but the entire biochemical process of CO_2_ assimilation, including mesophyll and lower level (i.e., cell wall, plasma membrane, cytosol) conductance. Additionally, differences in nutrient acquisition and use patterns among scrub *R. mangle* trees growing at island edges vs. centers affect leaf-nutrient status and photosynthetic potential.

The findings from this study indicate that it is the interaction of inundation stress with mangrove-island micro-elevational habitat in the flooded scrub mangroves of the southeastern Florida Everglades that principally alters tree water and nutrient-use dynamics, which appear to cascade to affect leaf gas exchange rates through their effects on *g_sw_*. Reductions in *A_net_* interact with the salinity of the water that inundates scrub *R. mangle* trees, in theory, because *g_sw_* rates are low and primarily respond to water loss from leaves rather than carbon gain (see supplemental section on *R. mangle* CO_2_ assimilation and stomatal behavior). In our field measurements, however, we found that prolonged inundation more than porewater salinity drives reduction in *A_net_* because the hydrological regime at Taylor River is characterized by long hydroperiods and minor fluctuations in salinity throughout the year (Figure S2). At the forest level, such physiological differences in scrub mangrove functioning with habitat and hydrological environment can help inform ecosystem carbon cycle models and mangrove forest responses to SLR and saltwater intrusion.

## ACKNOWLEDGEMENTS

This research was supported by the Department of Interior – National Park Service (Cooperative Agreement #P16AC00032) and the Florida Coastal Everglades Long-Term Ecological Research (FCE-LTER) program funded by the National Science Foundation (Grant #DEB-1832229). We thank Mandy Kuhn, Austin Pezoldt, and Ralph Diaz-Hung for field assistance. We thank the Everglades National Park for granting research permits and the Florida Bay Interagency Science Center-Everglades National Park (FBISC-ENP) for logistic support during the study. We acknowledge Dr. Zoran Bursac from the Center for Statistical Consulting and Collaboration at FIU for statistical advice. Finally, we thank Prof. Ernesto Medina and two anonymous reviewers for constructive comments on an earlier version of this manuscript. This is contribution number 1040 from the Southeast Environmental Research Center in the Institute of Environment at Florida International University.

## CONFLICTS OF INTERST

None declared

## Notes

https://doi.org/10.6073/pasta/27f6332609eb1ef6d398c7855855f2e3

### Competing Interest Statement

The authors have declared no competing interest.

### Summary of Updates

This is the revised submission (after proofing, ACCEPTED IN Tree Physiology) doi: 10.1093/treephys/tpab151

## REFERENCES

Abiy AZ, Melesse AM, Abtew W, Whitman D (2019) Rainfall trend and variability in Southeast Florida: Implications for freshwater availability in the Everglades. PLoS One 14:e0212008.

Aerts R, Chapin FS. (2002) The mineral nutrition of wild plants revisited: a re-evaluation of processes and patterns. Adv Ecol Res 30:1–67.

Alongi DM (2008) Mangrove forests: Resilience, protection from tsunamis, and responses to global climate change. Estuarine Coastal Shelf Sci 76:1–13.

Alongi DM, Boto KG, Robertson AI (1992) Nitrogen and phosphorus cycles. In: Robertson AI, Alongi DM (eds) Tropical Mangrove Ecosystems. American Geophysical Union, Washington, DC.

Ball MC (2009) Photosynthesis in mangroves. Wetlands Australia 6:12–22.

Ball MC (1988) Ecophysiology of mangroves. Trees-Struct Funct 2:129–142.

Ball MC (1996) Comparative ecophysiology of mangrove forest and tropical lowland moist rainforest. In: Mulkey SS, Chazdon RL, Smith AP (eds) Tropical forest plant ecophysiology Springer, Boston, Massachusetts, pp 461–496.

Barr JG, Fuentes JD, Engel V, Zieman JC (2009) Physiological responses of red mangroves to the climate in the Florida Everglades. J Geophys Res Biogeosci 114:G02008.

Bates D, Mächler M, Bolker B, Walker S. (2015) Fitting linear fixed-effects models using lme4. J Stat Softw 67:1–48.

Bjorkman O, Demmig B, Andrews TJ (1988) Mangrove photosynthesis: response to high-irradiance stress. Funct Plant Bio 15:43–61.

Cardona-Olarte P, Krauss KW, Twilley RR (2013) Leaf gas exchange and nutrient use efficiency help explain the distribution of two Neotropical mangroves under contrasting flooding and salinity. Int J For Res 2013:1–10. doi:10.1155/2013/524625

Cardona-Olarte P, Twilley RR, Krauss KW, Rivera-Monroy VH (2006) Responses of neotropical mangrove seedlings grown in monoculture and mixed culture under treatments of hydroperiod and salinity. Hydrobiologia 569:325–341.

Castañeda-Moya E, Twilley RR, Rivera-Monroy VH, Marx BD, Coronado-Molina C, Ewe SML (2011) Patterns of root dynamics in mangrove forests along environmental gradients in the Florida Coastal Everglades, USA. Ecosystems 14:1178–1195.

Castañeda-Moya E, Twilley RR, Rivera-Monroy VH (2013) Allocation of biomass and net primary productivity of mangrove forests along environmental gradients in the Florida Coastal Everglades, USA. For Ecol Manage 307:226–241.

Chambers LG, Davis SE, Troxler T, Boyer JN, Downey-Wall A, Scinto LJ (2014) Biogeochemical effects of simulated sea level rise on carbon loss in an Everglades mangrove peat soil. Hydrobiologia 726:195–211.

Chapin FS, Bloom AJ, Field CB, Waring RH (1987) Plant responses to multiple environmental factors. BioScience 37:49–57.

Chapin FS, Moilanen L (1991) Nutritional controls over nitrogen and phosphorus resorption from Alaskan birch leaves. Ecology 72:709–715.

Cheeseman J, Herendeen L, Cheeseman A, Clough B (1997) Photosynthesis and photoprotection in mangroves under field conditions. Plant Cell Environ 20:579–588.

Cheeseman J, Lovelock C (2004) Photosynthetic characteristics of dwarf and fringe Rhizophora mangle L. in a Belizean mangrove. Plant Cell Environ 27:769–780.

Chen R, Twilley RR (1999) A simulation model of organic matter and nutrient accumulation in mangrove wetland soils. Biogeochemistry. 44:93–118.

Childers DL (2006) A synthesis of long-term research by the Florida Coastal Everglades LTER Program. Hydrobiologia. 569:531–544.

Cintron G, Lugo AE, Douglas D, Pool J, Morris G (1978) Mangroves of Arid Environments in Puerto Rico and Adjacent Islands. Biotropica 10:110–121.

Clough B, Sim R (1989) Changes in gas exchange characteristics and water use efficiency of mangroves in response to salinity and vapour pressure deficit. Oecologia 79:38–44.

Cornelissen JH, Lavorel S, Garnier E, Díaz S, Buchmann N, Gurvich DE, Reich PB, Ter Steege H, Morgan HD, Van Der Heijden MG, Pausas JG (2003) A handbook of protocols for standardised and easy measurement of plant functional traits worldwide. Aust J Bot 51:335–80.

Craine JM, Brooksheir ENJ, Cramer MD, Hasselquist NJ, Koba K, Marin-Spiotta E, Wang L (2015) Ecological interpreations of nitrogen isotope ratios of terrestrial plans and soils Plant Soil 396:1–26.

Duever MJ, Meeder JF, Meeder LC, McCollom JM (1994) The climate of South Florida and its role in shaping the Everglades ecosystem. In: Davis SM, Ogden JC (eds) Everglades: The Ecosystem and Its Restoration. St. Lucie Press, Delray Beach, Florida, pp 225–248.

Ellison AM, Farnsworth EJ (1997) Simulated sea level change alters anatomy, physiology, growth, and reproduction of red mangrove (*Rhizophora mangle* L.). Oecologia 112:435–446.

Ewe SM, Gaiser EE, Childers DL, Iwaniec D, Rivera-Monroy VH, Twilley RR (2006) Spatial and temporal patterns of aboveground net primary productivity (ANPP) along two freshwater-estuarine transects in the Florida Coastal Everglades. Hydrobiologia 569:459–474.

Ewe SM, Sternberg LSL, Childers DL. 2007. Seasonal plant water uptake patterns in the saline southeast Everglades ecotone. Oecologia 152: 607–616.

Farnsworth EJ, Ellison AM (1996) Sun-shade adaptability of the red mangrove, *Rhizophora mangle* (Rhizophoraceae): Changes through ontogeny at several levels of biological organization. Amer J Bot 83:1131–1143.

Feller IC (1995) Effects of nutrient enrichment on growth and herbivory of dwarf red mangrove (Rhizophora mangle). Ecol Monogr 65:477–505.

Feller IC, Whigham DF, O’Neil JP, McKee KL (1999) Effects of nutrient enrichement on within-stand cycling in a mangrove forest. Ecology. 80:2193–2205.

Feller IC, McKee KL, Whigham DF, O’Neill JP (2003a) Nitrogen vs. phosphorus limitation across an ecotonal gradient in a mangrove forest. Biogeochemistry 62:145–175.

Feller IC, Whigham DF, McKee KL, Lovelock CE (2003b) Nitrogen limitation of growth and nutrient dynamics in a disturbed mangrove forest, Indian River Lagoon, Florida. Oecologia 134:405–414.

Field CD (1984) Ions in mangroves. In: Teas HJ (ed) Physiology and management of mangroves. Springer, Dordrecht, Netherlands, pp 43–48.

Field CD (1995) Impact of expected climate change on mangroves. Hydrobiologia 295:75–81.

Fogel M, Wooller M, Cheeseman J, Smallwood B, Roberts Q, Romero I, Jacobsen Meyers M (2008) Unusually negative nitrogen isotopic compositions (δ^15^N) of mangroves and lichens in an oligotrophic, microbially-influenced ecosystem. Biogeosci Discuss 5:1693–1704.

Fry B, Bern A, Ross M, Meeder J (2000) δ^15^N studies of nitrogen use by the red mangrove, Rhizophora mangle L. in south Florida. Estuarine Coastal Shelf Sci 50:291–296.

Garten CTJ (1993) Variation in foliar ^15^N abundance and the availability of soil nitrogen on Walker Branch watershed. Ecology 74:2098–2113.

Gleeson SK, Tilman D (1992) Plant allocation and the multiple limitation hypothesis. Am Nat 139:1322–1343.

Golley F, Odum HT, Wilson RF (1962) The structure and metabolism of a Puerto Rican red mangrove forest in May. Ecology 43:9–19.

He B, Lai T, Fan H, Wang W, Zheng H (2007) Comparison of flooding-tolerance in four mangrove species in a diurnal tidal zone in the Beibu Gulf. Estuarine Coastal Shelf Sci 74:254–262.

He D, Rivera-Monroy VH, Jaffé R, Zhao X (2020) Mangrove leaf species-specific isotopic signatures along a salinity and phosphorus soil fertility gradients in a subtropical estuary. Estuarine Coastal Shelf Sci 248:106768.

Högberg P (1997) ^15^N natural abundance in soil-plant systems. New Phytol 137:179–203.

Hoppe-Speer SC, Adams JB, Rajkaran A, Bailey D (2011) The response of the red mangrove *Rhizophora mucronata* Lam. to salinity and inundation in South Africa. Aquat Bot 95:71–76.

Hughes CE, Binning P, Willgoose GR (1998) Characterisation of the hydrology of an estuarine wetland. J Hydrol 211:34–49.

Koch MS, Snedaker SC (1997) Factors influencing *Rhizophora mangle* L. seedling development in Everglades carbonate soils. Aquat Bot 59:87–98.

Kozlowski TT (1997) Responses of woody plants to flooding and salinity. Tree Physiol 17:490–490.

Krauss KW, McKee KL, Lovelock CE, Cahoon DR, Saintilan N, Reef R, Chen L (2014) How mangrove forests adjust to rising sea level. New Phytol 202:19–34.

Krauss KW, Twilley RR, Doyle TW, Gardiner ES (2006) Leaf gas exchange characteristics of three neotropical mangrove species in response to varying hydroperiod. Tree Physiol 26:959–968.

Kuznetsova A, Brockhoff PB, Christensen RHB (2017) lmerTest Package: Tests in Linear Mixed Effects Models. J Stat Softw 82:13.

Lambers H, Chapin FS, Pons TL (eds) (2008) Plant physiological ecology. Springer, New York, New York.

Lamers LPM, Govers LL, Janssen ICJM, Geurts JJM, Van der Welle MEW, Van Katwijk MM, Van del Heide T, Roelofs JGM, Smolders AJP (2013) Sulfide as a soil phytotoxin - a review. Front Plant Sci 4:1–14.

Lawton JR, Todd ANN, Naidoo DK (1981) Preliminary inverstigations into the structure of the roots of mangroves, *Avicennia marina* and *Brugeria gymnorrhiza*, in relation to ion uptake. New Phytol 88:713–722.

Lin G, Sternberg LSL (1992) Comparative study of water uptake and photosynthetic gas exchange between scrub and fringe red mangroves, *Rhizophora mangle* L. Oecologia 90:399–403.

Lopes D, Tognella M, Falqueto A, Soares M (2019) Salinity variation effects on photosynthetic responses of the mangrove species *Rhizophora mangle* L. growing in natural habitats. Photosynthetica 57:1142–1155.

Loveless CM (1959) A study of the vegetation in the Florida Everglades. Ecology 40:2–9.

Lovelock CE (2008) Soil respiration and belowground carbon allocation in mangrove forests. Ecosystems 11:342–354.

Lovelock CE, Feller I (2003) Photosynthetic performance and resource utilization of two mangrove species coexisting in hypersaline scrub forest. Oecologia 134:455–462.

Lovelock CE, Feller IC, McKee KL, Engelbrecht BM, Ball MC (2004) The effect of nutrient enrichment on growth, photosynthesis and hydraulic conductance of dwarf mangroves in Panama. Funct Ecol 18:25–33.

Lovelock CE, Atwood T, Baldock J, Duarte CM, Hickey S, Lavery P, Masque P, Macreadie PI, Ricart AM, Serrano O, Steven A (2017) Assesing the risk of carbon dioxide emissions from blue carbon ecosystems. Front Ecol Environ 15:257–265.

Lüdecke D (2018) sjPlot: Data visualization for statistics in social science. R package version 2. https://cran.r-project.org/web/packages/sjPlot/

Lugo AE, Medina E, Cuevas E, Cintrón G, Nieves ENL, Novelli YS (2007) Ecophysiology of a mangrove forest in Jobos Bay, Puerto Rico. Carib J Sci 43:200–219.

Lugo AE, Snedaker SC (1974) The ecology of mangroves. Ann Rev Ecol System 5:39–64.

Mancera-Pineda JE, Twilley RR, Rivera-Monroy VH (2009) Carbon (δ^13^C) and nitrogen (δ ^15^N) isotopic discrimination in mangroves in Florida coastal Everglades as a function of enviromnetal stress. Contrib Mar Sci 38:109–129.

Marshall JD, Brooks JR, Lajtha K (2007) Stable isotopes in ecology and environmental science. In: Michener R, Lajtha K (eds) Sources of variation in the stable isotopic composition of plants. Blackwell, Singapore, pp 22–60.

McKee KL (1993) Soil physicochemical patterns and mangrove species distribution-reciprocal effects? J Ecol 81:477–487.

McKee KL (2011) Biophysical controls on accretion and elevation change in Caribbean mangrove ecosystems. Estuarine Coastal Shelf Sci 91:475–483.

McKee KL, Cahoon DR, Feller IC (2007) Caribbean mangroves adjust to rising sea level through biotic controls on change in soil elevation. Global Ecol Biogeogr 16:545–556.

McKee KL, Feller IC, Popp M, Wanek W (2002) Mangrove isotopic (δ^15^N and δ^13^C) fractionation across a nitrogen vs. phosphorus limitation gradient. Ecology 83:1065–1075.

Mcleod E, Chmura GL, Bouillon S, Salm R, Bjork M, Duarte CM, Lovelock CE, Schlesinger WH, Silliman BR (2011) A blueprint for blue carbon: Toward an improved understanding of the role of vegetated coastal habitats in sequestering CO_2_. Front Ecol Environ 9:552–560.

Medina E (1999) Mangrove physiology: the challenge of salt, heat, and light stress under recurrent flooding. In: Yañez-Arancibia A, Lara-Domínguez L (eds) Ecosistemas de manglar en América tropical. Instituto de Ecología A.C. México, UICN/ORMA, Costa Rica, NOAA/NMFS Silver Spring, MD USA. pp 109–126.

Medina E, Cuevas E, Lugo AE (2010) Nutrient relations of dwarf *Rhizophora mangle* L. mangroves on peat in eastern Puerto Rico. Plant ecolog 207:13–24.

Medina E, Francisco M (1997) Osmolality and δ^13^C of leaf tissues of mangrove species from environments of contrasting rainfall and salinity. Estuarine Coastal Shelf Sci 45:337–344.

Medina E, Francisco M, Quilice A (2008) Isotopic signatures and nutrient relations of plants inhabiting brackish wetlands in the northeastern coastal plain of Venezuela. Wetlands Ecol Manage 16:51.

Mendelssohn I, McKee K (2000) Saltmarshes and mangroves. In: Barbour MG, Billings WD (eds) North American terrestrial vegetation. Cambridge University Press, New York, NY, pp 501–536.

Michot B, Meselhe EA, Rivera-Monroy VH, Coronado-Molina C, Twilley RR (2011) A tidal creek water budget: estimation of groundwater discharge and overland flow using hydrologic modeling in the southern Everglades. Estuarine Coastal Shelf Sci 93:438–448.

Murdiyarso D, Purbopuspito J, Kauffman JB, Warren MW, Sasmito SD, Donato DC, Manuri S, Krisnawati H, Taberima S, Kurnianto S (2015) The potential of Indonesian mangrove forests for global climate change mitigation. Nat Clim Change 5:1089–1092.

Naidoo G (1983) Effects of flooding on leaf water potential and stomatal resistance in *Bruguiera* gymnorrhiza (L.) LAM. New Phytol 93:369–373.

Neubauer SC, Franklin RB, Berrier DJ (2013) Saltwater intrusion into tidal freshwater marshes alters the biogeochemical processing of organic carbon. Biogeosciences Discuss 10:10685–10720.

Nickerson NH, Thibodeau FR (1985) Association between porewater sulfide concentrations and distribution of mangroves. Biogeochemistry 1:183–192.

O’Leary MH (1988) Carbon isotopes in photosynthesis. BioScience. 38:328–336.

Parida AK, Jha B (2010) Salt tolerance mechanisms in mangroves: a review. Trees-Struct Funct 24:199–217.

Pezeshki S, DeLaune R, Patrick Jr W (1990a) Flooding and saltwater intrusion: Potential effects on survival and productivity of wetland forests along the U.S. Gulf Coast. For Ecol Manage 33:287–301.

Pezeshki S, DeLaune R, Patrick Jr W (1990b) Differential response of selected mangroves to soil flooding and salinity: gas exchange and biomass partitioning. Can J For Res 20:869–874.

Pezeshki SR, DeLaune RD (2012) Soil oxidation-reduction in wetlands and its impact on plant functioning. Biology 1:196–221.

Potts M (1979) Nitrogen fixation (acetylene reduction) associated with communities of heterocystous and non-heterocystous blue-green algae in mangrove forests of Sinai. Oecologia 39:359–373.

Pugnaire FI, Chapin FS (1993) Controls over nutrient resorption from leaves of evergreen Mediterranean species. Ecology 4:124–129.

Price RM, PK Swart, Fourqurean JW (2006) Coastal groundwater discharge - an additional source of phosphorus for the oligotrophic wetlands of the Everglades. Hydrobiologia 569: 23–36.

R Core Team (2018) R: a language and environment for statiscal computing. R foundation for staistical computing. Vienna, Austria. www.r-project.org

Reef R, Lovelock CE (2015) Regulation of water balance in mangroves. Ann Bot 115:385–395.

Reis CRG, Nardoto GB, Oliveira RS (2017a) Global overview on nitrogen dynamics in mangroves and consequences of increasing nitrogen availability for these systems. Plant Soil 410:1–19.

Reis CRG, Nardoto GB, Rochelle ALC, Vieira SA, Oliveira RS (2017b) Nitrogen dynamics in subtropical fringe and basin mangrove forests inferred from stable isotopes. Oecologia 183:841–848.

Robinson D (2001) δ^15^N as an integrator of the nitrogen cycle. Trends Ecol Evol 16:153–162.

Ross MS, Meeder JF, Sah JP, Ruiz PL, Telesnicki GJ (2000) The southeast saline Everglades revisited: 50 years of coastal vegetation change. J Veg Sci 11:101–112.

Rovai AS, Twilley RR, Castañeda-Moya E, Riul P, Cifuentes-Lara M, Manrow-Villalobos M, Horta PA, Simonassi JC, Fonseca AL, Pagliosa PR (2018) Global controls of carbon storage in mangrove soils. Nat Clim Change 8:534–538.

Schneider CA, Rasband WS, Eliceiri KW (2012) NIH Image to ImageJ: 25 years of image analysis. Nat Methods 9:671–675.

Scholander P (1968) How mangroves desalinate seawater. Physiol Plant 21:251–261.

Scholander P, Hammel H, Hemmingsen E, Garey W (1962) Salt balance in mangroves. Plant Physiol 37:722.

Sobrado M (2000) Relation of water transport to leaf gas exchange properties in three mangrove species. Tree-Struct Funct 14:258–262.

Sutula M, Day JW, Cable JE, Rudnick D (2001) Hydrological and nutrient budgets of freshwater and estuarine wetlands of Taylor Slough in southern Everglades, Florida (U.S.A.). Biogeochemistry 56:287–310.

Tully K, Gedan K, Epanchin-Niell R, Strong A, Bernhardt ES, BenDor T, Mitchell M, Kominoski J, Jordan TE, Neubauer SC, Weston NB (2019) The invisible flood: The chemistry, ecology, and social implications of coastal saltwater intrusion. BioScience 69:368–378.

Tomlinson PB (2016) The botany of mangroves. Cambridge University Press, Cambridge, Massachusetts.

Troxler TG (2007) Patterns of phosphorus, nitrogen and δ^15^N along a peat development gradient in a coastal mire, Panama. J Trop Ecology 23:683–691.

Twilley RR, Castañeda-Moya E, Rivera-Monroy VH, Rovai A (2017) Productivity and carbon dynamics in mangrove wetlands. In: Rivera-Monroy VH, Lee SY, Kristensen E, Twilley RR (eds) Mangrove ecosystems: A global biogeographic perspective. Springer International, Cham, Switzerland, pp 113–162.

Twilley RR, Lugo AE, Patterson-Zucca C (1986) Litter production and turnover in basin mangrove forests in southwest Florida. Ecology 67:670–683.

Twilley RR, Rivera-Monroy V (2005) Developing performance measures of mangrove wetlands using simulation models of hydrology, nutrient biogeochemistry, and community dynamics. J Coast Res 40:79–93.

Twilley RR, Rivera-Monroy V (2009) Ecogeomorphic models of nutrient biogeochemistry for mangrove wetlands. In: Coastal Wetlands: An integrated ecosystem approach (eds) Perillo GME, Wolanski E, Cahoon DR and Brinson MM. Elsevier, Amsterdam, Netherlands, pp 641–683.

Twilley RR, Rivera-Monroy V, Rovai AS, Castañeda-Moya E, Davis SE, III. (2019) Mangrove biogeochemistry at local to global scales using ecogeomorphic approaches. In: Perillo GME, Wolanski E, Cahoon DR, Hopkinson CS (eds) Coastal Wetlands: An Integrated Ecosystem Approach.Elsevier, Amsterdam, Netherlands, pp 717–785.

Twilley RR, Rivera-Monroy VH, Chen R, Botero L (1998) Adapting and ecological mangrove model to simulate trajectories in restoration ecology. Mar. Poll. Bull. 37:404–419.

Uncles R, Gong W-K, Ong J-E (1992) Intratidal fluctuations in stratification within a mangrove estuary. In: Martens E, Jaccarini V (eds) The Ecology of Mangrove and Related Ecosystems. Springer, Dordrecht, Germany, pp 163–171.

US EPA (1983) Methods for chemical analysis of water and wastes. United States Environmental Protection Agency Office of Research and Development, Washington, DC.

Vitousek P (1982) Nutrient cycling and nutrient use efficiency. Am Nat 119:553–572.

Wanless H (1998) Mangroves, hurricanes, and sea level rise. South Florida Study Group, The Conservancy, Naples, Florida

Werner A, Stelzer R (1990) Physiological responses of the mangrove *Rhizophora mangle* grown in the absence and presence of NaCl. Plant Cell Environ 13:243–255.

Wolanski E (1992) Hydrodynamics of mangrove swamps and their coastal waters. Hydrobiologia 247:141–161.

Wooller M, Smallwood B, Scharler U, Jacobson M, Fogel M (2003) A taphonomic study of δ13C and δ15N values in Rhizophora mangle leaves for a multi-proxy approach to mangrove palaeoecology. Org Geochem 34:1259–1275.

Yu M, Rivera-Ocasio E, Heartsill-Scalley T, Davila-Casanova D, Rios-López N, Gao Q. (2019) Landscape-level consequences of rising sea-level on coastal wetlands: saltwater intrusion drives displacement and mortality in the twenty-first century. Wetlands 39:1343–1355.

