## supplemental for "Water levels primarily drive variation in photosynthesis and nutrient use of scrub Red Mangroves in the southeastern Florida Everglades" (from another version of this preprint: version 4, 2021-10-15)

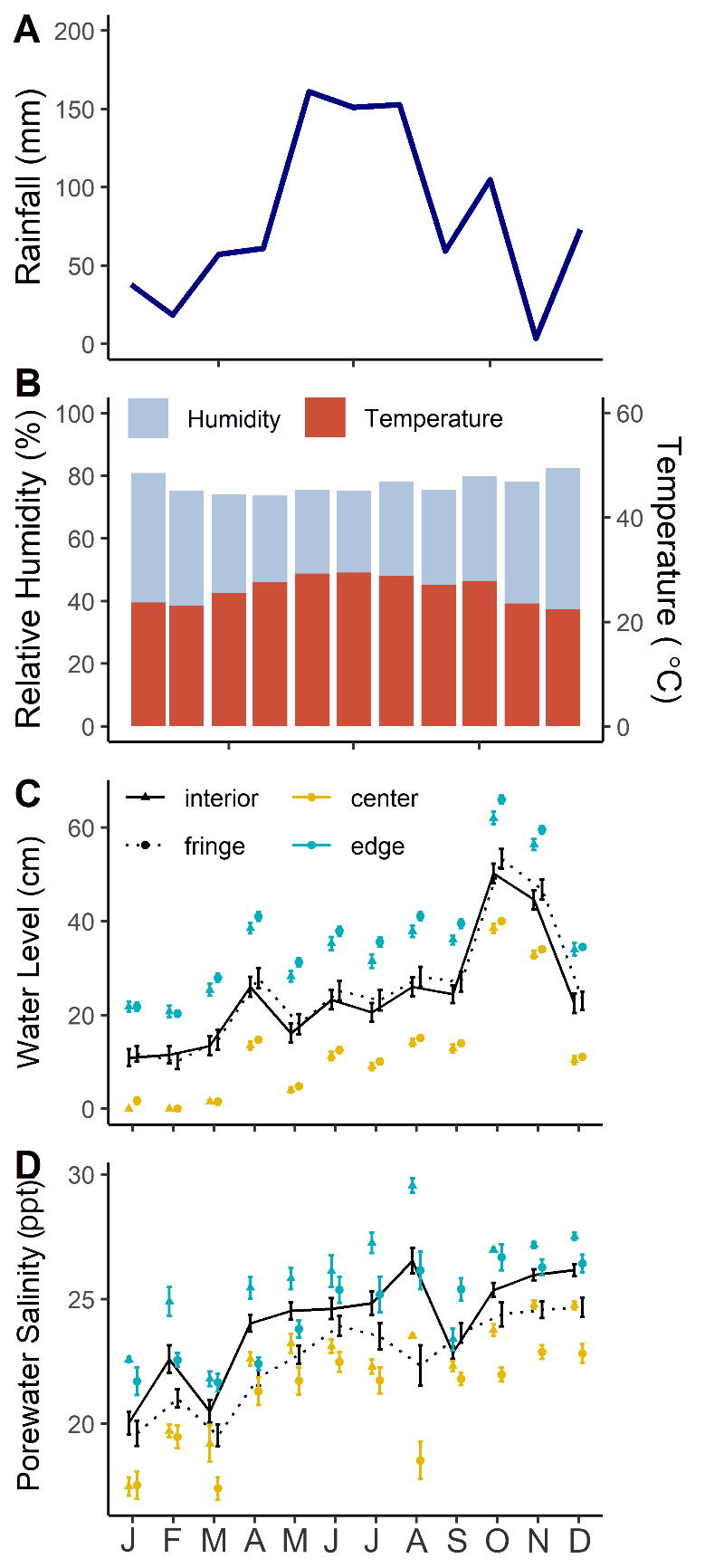

**Figure S1.** Environmental parameters for TS/Ph-7 for the 2019 calendar year. Cumulative monthly rainfall **(A)**, monthly average relative humidity and air temperature **(B)**, average (± 1 SE) water level **(C),** and average (± 1 SE) porewater salinity **(D)** measured at mangrove island habitats at the study site. Scrub mangroves from fringe and interior zones sampled during the study are shown in dotted and solid lines, respectively, whereas mangrove edge, and center habitats are differentiated by color (teal and gold, respectively). Rainfall data **(A)** were obtained from a USGS-maintained rain gauge in Taylor River, which is part of the Everglades Depth Estimation Network (Skinner et al. 2009). Data for **B** were collected from an eddy covariance tower located at the study site. Water level **(C)** and porewater salinity **(D)** were measured for each island, at both the edge and center habitats, during monthly mangrove leaf photosynthesis measurements.

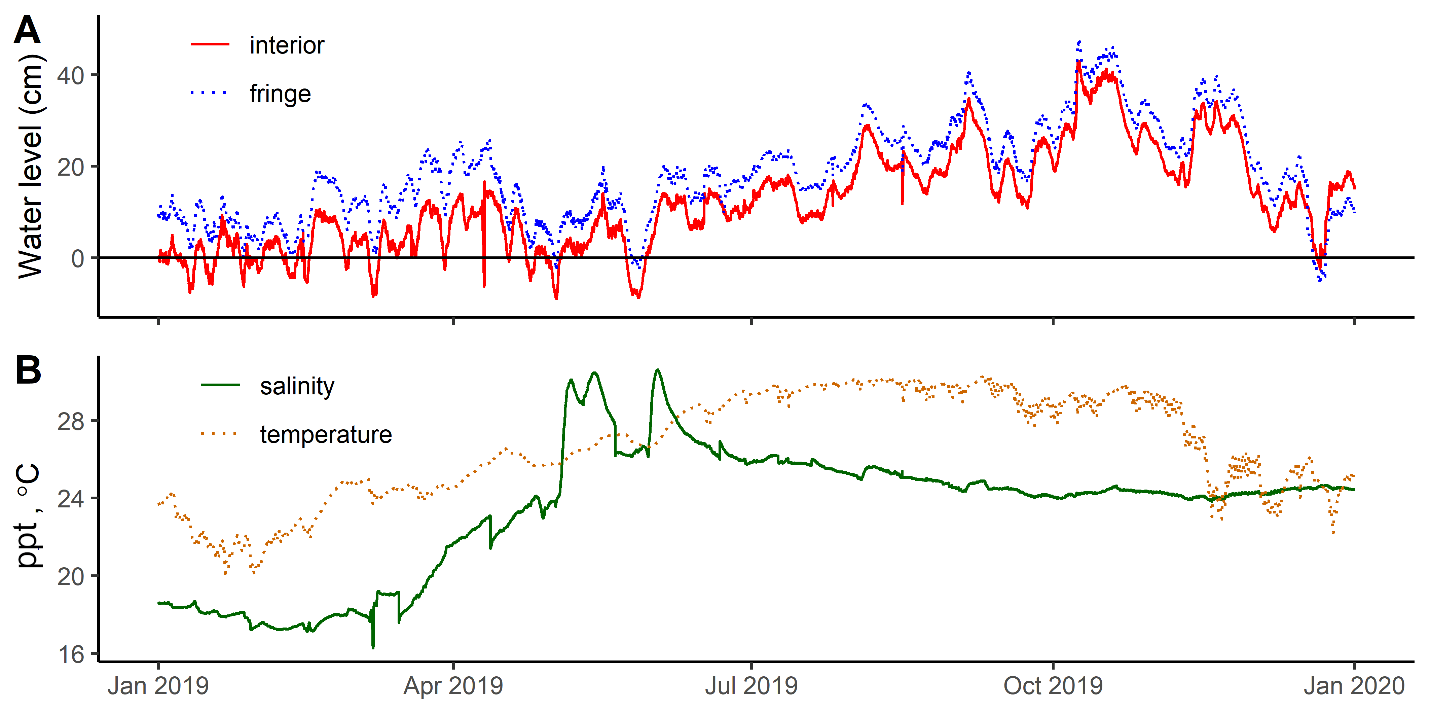

**Figure S2**: **(A)** Hydrographs in scrub mangrove zones (fringe and interior) at TS/Ph-7 from 2019. Data were collected using HOBO water level loggers (Interior zone; Onset Computer Corporation, MA) and Level TROLL (Fringe zone; In-Situ Inc., CO). The loggers were installed 100 m inland from the shoreline (interior) and 50 m from the edge of the mangrove zone (fringe). The zero mark is relative to the soil surface where each gauge was installed; water level data are not referenced to the North America Vertical Datum of 1988 (NAVD88). **(B)** Continuous measurements (1-h intervals, January-December 2019) of porewater salinity and temperature (30 cm depth) in the scrub fringe mangrove zone at TS/Ph-7 in southeastern Florida Everglades. Data were collected using a Level TROLL 200 Datalogger (In Situ Inc., Fort Collins, Colorado).

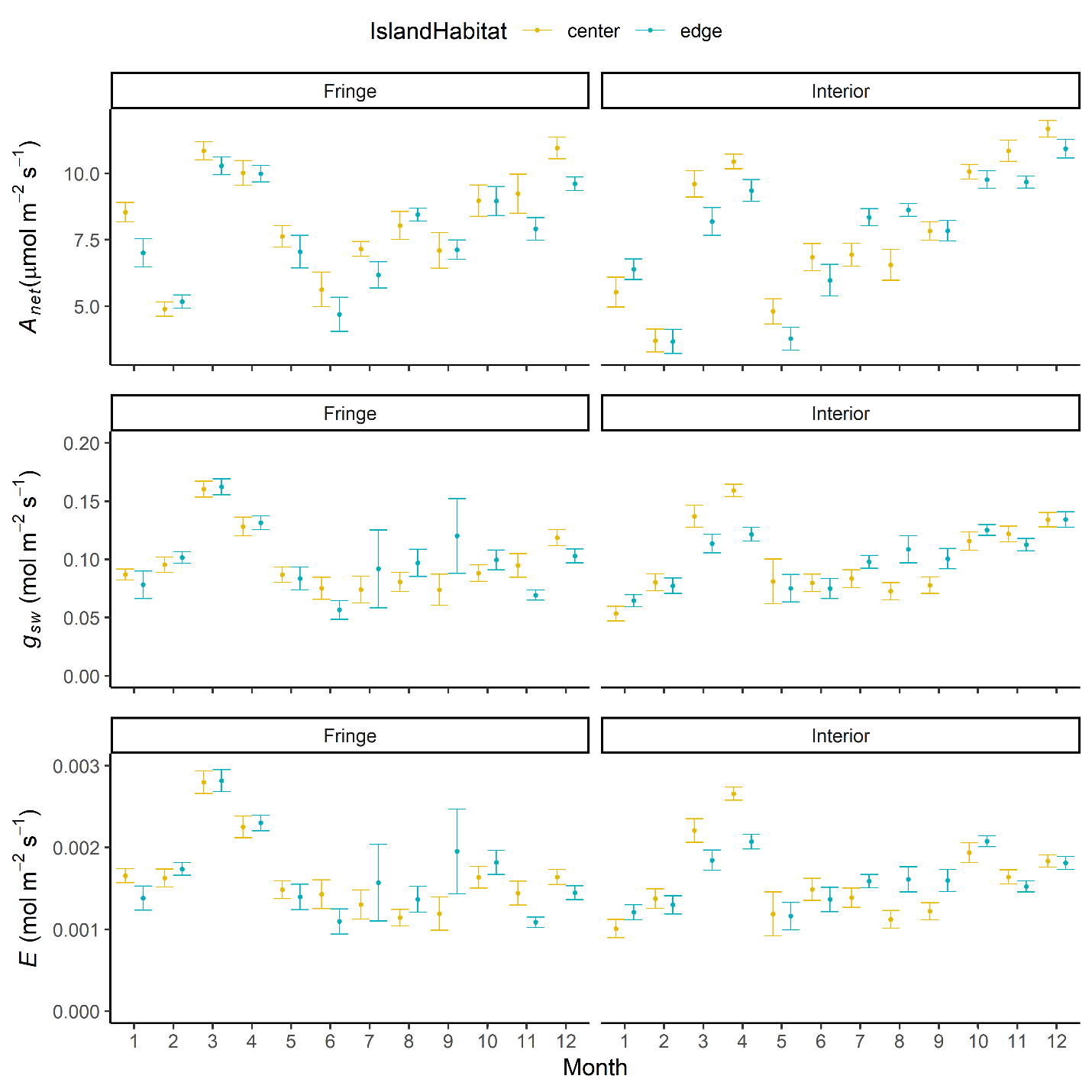

**Figure S3:** Temporal variation (means ± 1 SE) in leaf net CO_2_ assimilation (*A_net_*), stomatal conductance to water vapor (*g_sw_*), and transpiration (*E*) for the calendar year of 2019 for fringe and interior dwarf mangrove plots at TS/Ph-7 of the Florida Coastal Everglades LTER. Twenty leaves (i.e., half of the leaves, and 5 per island) were sampled per plot in each mangrove island habitat (center and edge).

**Figure S4 & Table S4:** Caterpillar plot and model summary table for the linear mixed-effect model of net photosynthetic rates (*A*). Fixed effect estimates, standard errors, confidence intervals, and associated probabilities for predictor terms are included in the model summary. A random effect for mangrove island was included in the model, therefore random effect variance (σ^2^) and intraclass correlation (ICC) are given. Overall model goodness of fit with (Conditional R^2^) and without (Marginal R^2^) random terms, AIC, deviance, and log likelihood are also shown.

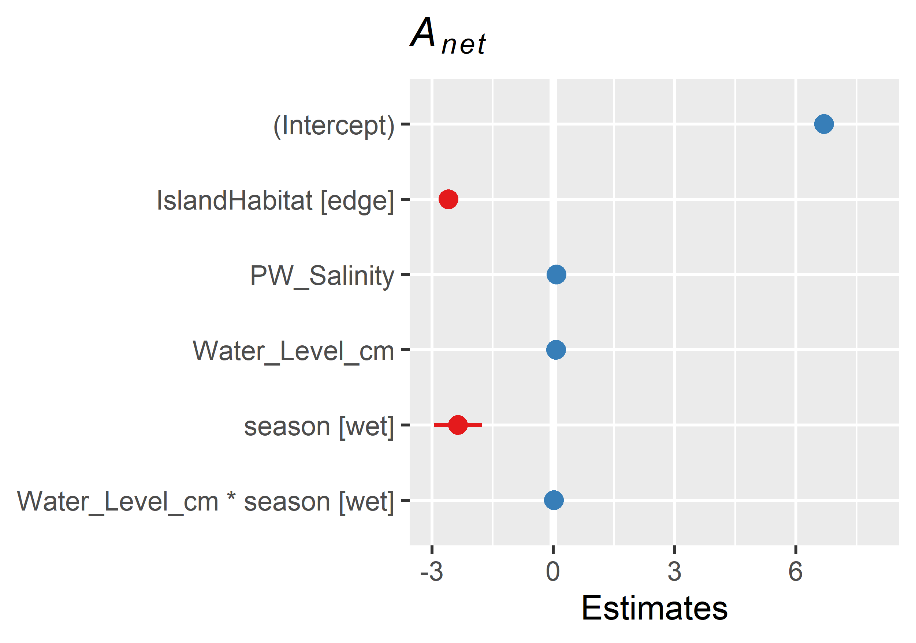

|  | ***A_net_*** | | | |
| --- | --- | --- | --- | --- |
| *Predictors* | *Estimates* | *std. Error* | *CI* | *p* |
| (Intercept) | 6.57 | 0.72 | 5.16 – 7.98 | **<0.001** |
| IslandHabitat [edge] | -2.65 | 0.23 | -3.11 – -2.20 | **<0.001** |
| PW_Salinity | 0.08 | 0.04 | 0.01 – 0.15 | **0.025** |
| Water_Level_cm | 0.07 | 0.01 | 0.06 – 0.09 | **<0.001** |
| season [wet] | -2.39 | 0.30 | -2.98 – -1.80 | **<0.001** |
| Water_Level_cm * season [wet] | 0.02 | 0.01 | -0.00 – 0.04 | 0.053 |
| **Random Effects** | | | | |
| σ^2^ | 6.07 | | | |
| τ_00_ _Island_ | 0.18 | | | |
| ICC | 0.03 | | | |
| N _Island_ | 8 | | | |
| Observations | 920 | | | |
| Marginal R^2^ / Conditional R^2^ | 0.219 / 0.242 | | | |
| Deviance | 4275.751 | | | |
| AIC | 4317.621 | | | |
| log-Likelihood | -2150.811 | | | |

**Figure S5 & Table S5:** Caterpillar plot and model summary table for the linear mixed-effect model for leaf stomatal conductance (*g_sw_*). Fixed effect estimates, standard errors, confidence intervals, and associated probabilities for predictor terms are included in the model summary. A random effect for mangrove island was included in the model, therefore random effect variance (σ^2^) and intraclass correlation (ICC) are given. Overall model goodness of fit with (Conditional R^2^) and without (Marginal R^2^) random terms, AIC, deviance and log likelihood are also shown.

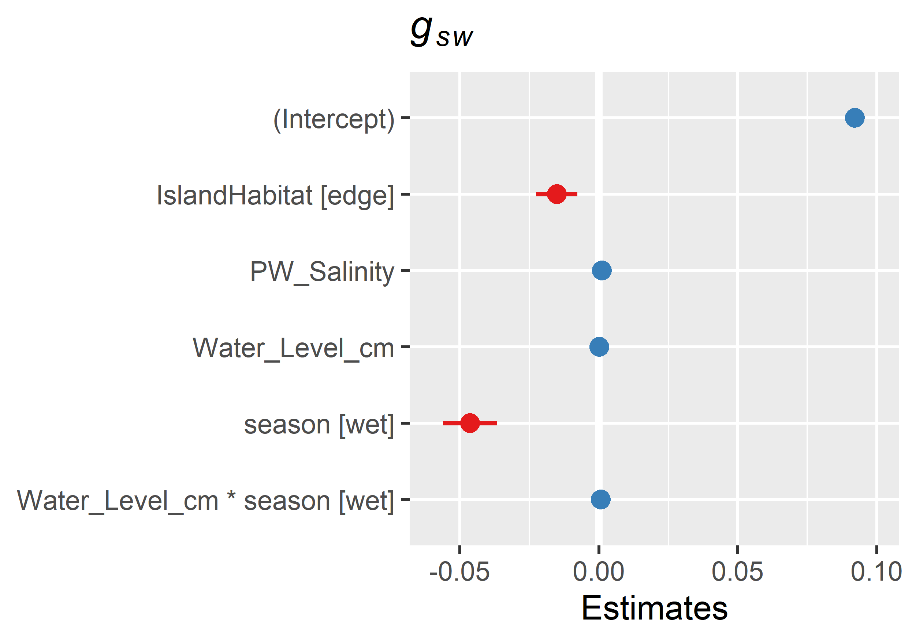

|  | ***g_sw_*** | | | |
| --- | --- | --- | --- | --- |
| *Predictors* | *Estimates* | *std. Error* | *CI* | *p* |
| (Intercept) | 0.09 | 0.01 | 0.07 – 0.12 | **<0.001** |
| IslandHabitat [edge] | -0.02 | 0.00 | -0.02 – -0.01 | **<0.001** |
| PW_Salinity | 0.00 | 0.00 | -0.00 – 0.00 | 0.069 |
| Water_Level_cm | 0.00 | 0.00 | -0.00 – 0.00 | 0.432 |
| season [wet] | -0.05 | 0.00 | -0.06 – -0.04 | **<0.001** |
| Water_Level_cm * season [wet] | 0.00 | 0.00 | 0.00 – 0.00 | **<0.001** |
| **Random Effects** | | | | |
| σ^2^ | 0.00 | | | |
| τ_00_ _Island_ | 0.00 | | | |
| ICC | 0.03 | | | |
| N _Island_ | 8 | | | |
| Observations | 920 | | | |
| Marginal R^2^ / Conditional R^2^ | 0.122 / 0.153 | | | |
| Deviance | -3286.781 | | | |
| AIC | -3195.746 | | | |
| log-Likelihood | 1605.873 | | | |

**Figure S6 & Table S6:** Caterpillar plot and model summary table for the linear mixed-effect model of the concentration of intracellular CO_2_ (*c_i_* Fixed effect estimates, standard errors, confidence intervals, and associated probabilities for predictor terms are included in the model summary. A random effect for mangrove island was included in the model, therefore random effect variance (σ^2^) and intraclass correlation (ICC) are given. Overall model goodness of fit with (Conditional R^2^) and without (Marginal R^2^) random terms, AIC, deviance and log likelihood are also shown.

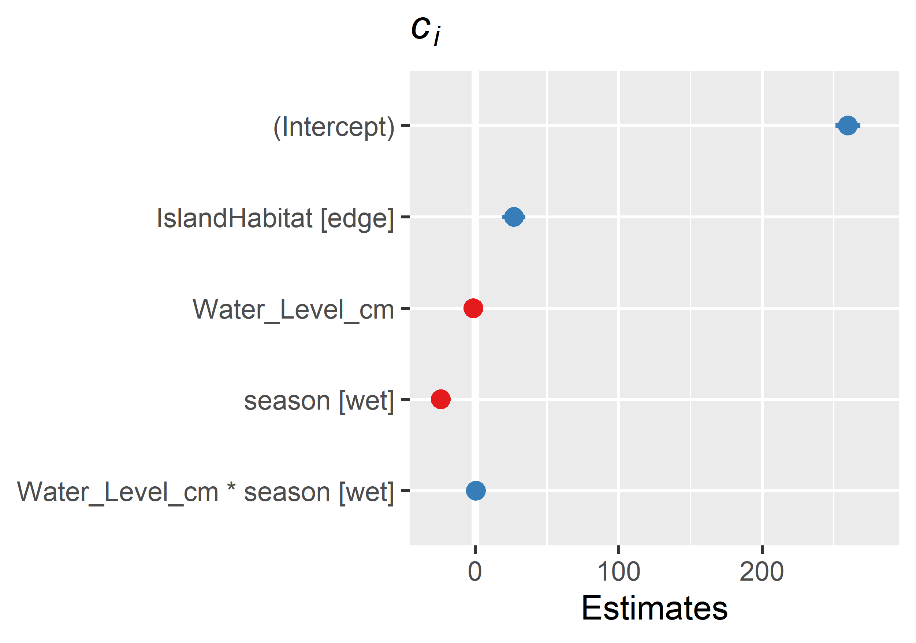

|  | ***c_i_*** | | | |
| --- | --- | --- | --- | --- |
| *Predictors* | *Estimates* | *std. Error* | *CI* | *p* |
| (Intercept) | 259.72 | 4.43 | 251.05 – 268.40 | **<0.001** |
| IslandHabitat [edge] | 27.07 | 4.13 | 18.98 – 35.16 | **<0.001** |
| Water_Level_cm | -1.15 | 0.14 | -1.43 – -0.87 | **<0.001** |
| season [wet] | -23.99 | 5.33 | -34.43 – -13.55 | **<0.001** |
| Water_Level_cm * season [wet] | 0.64 | 0.17 | 0.30 – 0.98 | **<0.001** |
| **Random Effects** | | | | |
| σ^2^ | 127.75 | | | |
| τ_00_ _Island_ | 65.87 | | | |
| ICC | 0.03 | | | |
| N _Island_ | 8 | | | |
| Observations | 920 | | | |
| Marginal R^2^ / Conditional R^2^ | 0.087 / 0.116 | | | |
| Deviance | 9655.152 | | | |
| AIC | 9661.042 | | | |
| log-Likelihood | -4823.521 | | | |

**Figure S7 & Table S7:** Caterpillar plot and model summary table for the linear mixed-effect model of photosynthetic water use efficiency (*wue*). Fixed effect estimates, standard errors, confidence intervals, and associated probabilities for predictor terms are included in the model summary. A random effect for mangrove island was included in the model, therefore random effect variance (σ^2^) and intraclass correlation (ICC) are given. Overall model goodness of fit with (Conditional R^2^) and without (Marginal R^2^) random terms, AIC, deviance and log likelihood are also shown.

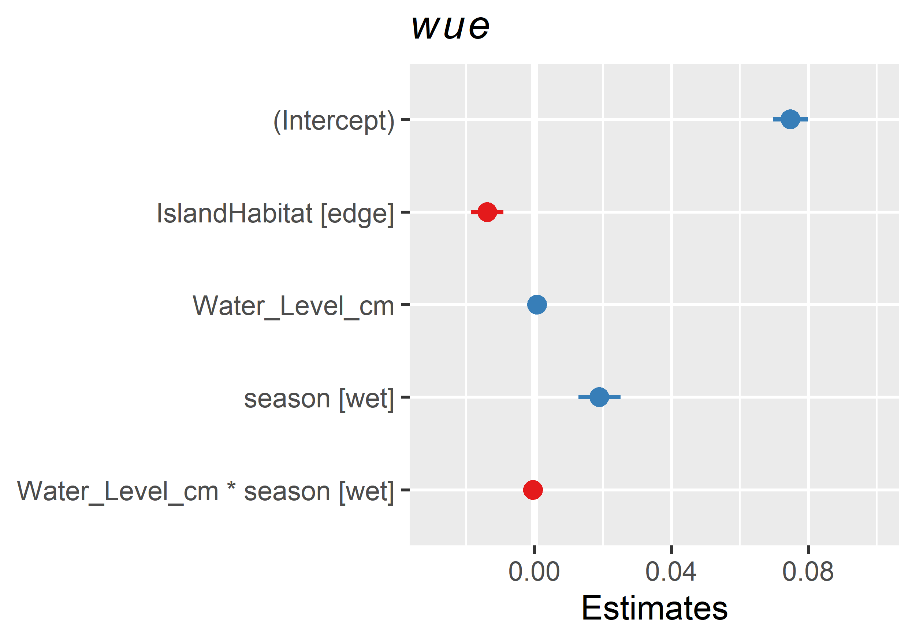

|  | ***wue*** | | | |
| --- | --- | --- | --- | --- |
| *Predictors* | *Estimates* | *std. Error* | *CI* | *p* |
| (Intercept) | 0.07 | 0.00 | 0.07 – 0.08 | **<0.001** |
| IslandHabitat [edge] | -0.01 | 0.00 | -0.02 – -0.01 | **<0.001** |
| Water_Level_cm | 0.00 | 0.00 | 0.00 – 0.00 | **<0.001** |
| season [wet] | 0.02 | 0.00 | 0.01 – 0.03 | **<0.001** |
| Water_Level_cm * season [wet] | -0.00 | 0.00 | -0.00 – -0.00 | **<0.001** |
| **Random Effects** | | | | |
| σ^2^ | 0.00 | | | |
| τ_00_ _Island_ | 0.00 | | | |
| ICC | 0.03 | | | |
| N _Island_ | 8 | | | |
| Observations | 920 | | | |
| Marginal R^2^ / Conditional R^2^ | 0.093 / 0.122 | | | |
| Deviance | -4066.233 | | | |
| AIC | -3985.972 | | | |
| log-Likelihood | 1999.986 | | | |

### Rhizophora mangle CO_2_ assimilation and stomatal behavior

The *A_net_-c_i_* relationship for *R. mangle* (Figure S8) and other salt-tolerant wetland plants warrants discussion because it results in unusual stomatal behavior (i.e., explains low rates of *g_sw_*, Ball 2009). To understand this relationship, we must realize how salinity affects *A_net_* in relation to *g_sw_.* Increasing salinity results in a downward translation of the *A_net_*-*c_i_* curve (i.e., lowers *A_max_*) and a decline in the initial slope of the *A_net_*-*c_i_* relationship (i.e., a proportional reduction in *A_net_* per µmol mol^-1^ increase in *c_i_*) (see Figure 3 in Ball 2009, taken from Ball and Farquhar 1984). Additionally, mangrove leaves almost always function at low *g_sw_* (i.e., typically <0.25 mol m^-2^ s^-1^ and “with an operational *c_i_* in the transition region of the A-ci curve,” *sensu* Ball 2009) with *c_i_* values of <200 µmol mol^-1^ (as reported by Ball 2009) to limit water loss relative to carbon gain and to maximize *wue.* Ball (2009) describes two hypothetical scenarios that typify changes in the *A_net_-c_i_* relationship for mangroves. The first assumes no change in the leaf's photosynthetic potential (i.e., no change in the *A_net_-c_i_* characteristics), which results in a decrease in *c_i_* and an increase in *wue* when stomates close at the expense of *A_net_*. The second scenario occurs when the photosynthetic potential of the leaf increases in concert with stomatal closure (i.e., a decrease in *g_sw_*), which results in a similar decrease in *c_i_* and an increase in *wue*, however at the expense of leaf nitrogen because of the increase in the photosynthetic machinery per unit leaf area. We observed differences in leaf nitrogen content that reflect differences in *A_net_* (Table 3, main text). Thus, variation in *c_i_* and *wue* reflect differences in the photosynthetic capacity of leaves among mangrove island habitats. Moreover, because mangrove leaves are particular in the degree to which they open their stomata in order to limit water loss, the differences in *g_sw_* between center and edge mangroves habitats are likely a function of inherent differences in the photosynthetic capacities of the leaves of mangrove trees in those respective habitats. Such differences might be attributable to differences in the acclimation of leaves to light intensity or because of inundation stress (e.g., lower rates of nutrient acquisition or sapflow), and may result from an interaction between light and flooding to optimally tune photosynthetic potential as a strategy for coping with inundation stress.

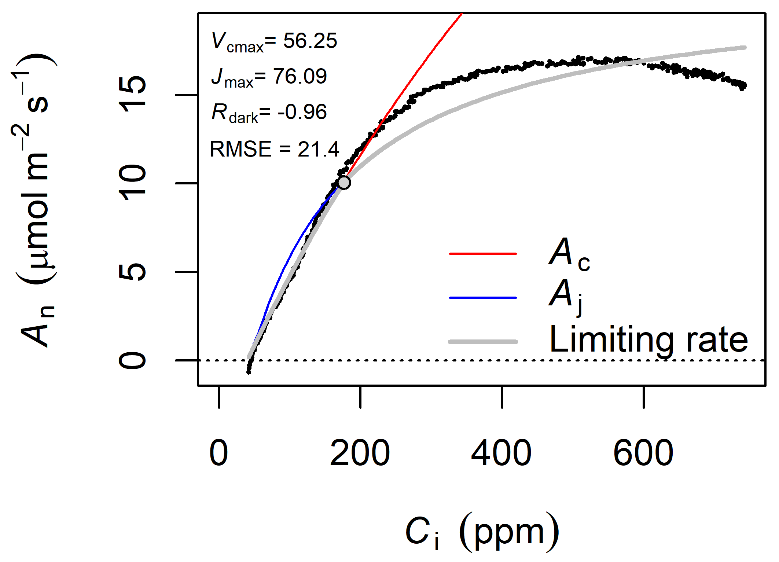

**Figure S9:** Photosynthetic-CO_2_ response (*A_n_*-*Ci*) curve for an *R. mangle* leaf collected from the scrub mangrove canopy at the TS/Ph-7 study site near the mouth of Taylor River in southeastern Everglades, Florida, USA. Measuring the CO_2_ response of photosynthesis on the severely nutrient-limited dwarf mangrove of TS/Ph-7 using the classical approach was difficult due to stomal closure. This curve was one of the best-measured responses, which was done using the RACiR technique (Stinziano et al. 2017). Stomatal closure begins to occur at about 600 µmols mol^-1^. Notably, our *V_cmax_* estimate is slightly lower than that reported by Barr et al. (2009, *V_cmax_* = 76.1 ± 23.4 sd), who measured the CO_2_ response of 29 leaves of tall *R. mangle* trees in Key Largo, FL.

**Works Cited in this Supplement**

Abiy AZ, Melesse AM, Abtew W, Whitman D. (2019) Rainfall trend and variability in Southeast Florida: Implications for freshwater availability in the Everglades. PloS ONE 14: e0212008.

Barr JG, Fuentes JD, Engel V, Zieman JC (2009) Physiological responses of red mangroves to the climate in the Florida Everglades. J Geophys Res Biogeosci 114:G02008.

Ball MC (2009) Photosynthesis in mangroves. Wetlands Australia 6:12-22.

Ball MC, Farquhar GD (1984) Photosynthetic and stomatal responses of two mangrove species, Aegiceras corniculatum and Avicennia marina, to long term salinity and humidity conditions. Plant Physiol 74:1-6.

Stinziano JR, Morgan PB, Lynch DJ, Saathoff AJ, McDermitt DK, Hanson DT (2017) The rapid A–Ci response: photosynthesis in the phenomic era. Plant Cell Environ 40:1256-1262.
